# MlaFEDB displays flippase activity to promote phospholipid transport towards the outer membrane of Gram-negative bacteria

**DOI:** 10.1101/2020.06.06.138008

**Authors:** Gareth W. Hughes, Pooja Sridhar, Stephanie A. Nestorow, Peter J. Wotherspoon, Benjamin F. Cooper, Timothy J. Knowles

## Abstract

MlaFEDB is a Gram-negative inner membrane protein complex involved in the inter membrane trafficking of phospholipids. Originally proposed to transport phospholipids in a retrograde direction, recent evidence suggests MlaFEDB may actually export phospholipids from the inner membrane to the periplasmic carrier protein, MlaC, potentially suggesting a role in either anterograde trafficking of phospholipids to the outer membrane or bidirectional phospholipid movement. MlaFEDB is part of the ABC transporter superfamily of proteins and has been shown to hydrolyse ATP through the cytoplasmic facing MlaF component. However, the movement of PLs from FEDB to MlaC has been shown to occur in an ATP independent fashion hence the role of ATP hydrolysis within this complex remains unclear. In this study we sought to elucidate the role of ATP and provide evidence to suggest MlaFEDB has flippase activity, utilising ATP hydrolysis to translocate phospholipids from the outer to the inner leaflet of the IM. We also show that in the absence of ATP MlaFEDB mediates the loading of MlaC with phospholipids directly from the inner leaflet only. Our data provides a novel role for MlaFEDB and presents a link between Mla driven phospholipid transport and ATP hydrolysis.

## Introduction

Gram-negative bacteria are characterised by an asymmetric outer membrane (OM) of phospholipids (PL) and lipopolysaccharides (LPS) separated from the inner bacterial membrane (IM) by the periplasmic space containing a thin peptidoglycan layer (Malinverni and Silhavy 2009). This envelope acts as a robust permeability barrier to a vast array of environmental substances that would otherwise threaten bacterial viability, with much of this barrier provided by the OM (Kamio and Nikaido 1976, Nikaido 2005, Ruiz et al. 2006). The integrity of this membrane is maintained by localisation of LPS to the outer leaflet and PL to the inner leaflet. It is well established that transport of LPS to the outer leaflet is conducted by the Lpt machinery (Tran et al. 2008, Chng et al. 2010, Okuda et al. 2016). The mechanisms by which PL are transported to the OM are less well understood. Multiple investigations, however, have begun to implicate the maintenance of lipid asymmetry (Mla) pathway in having a role in this by ferrying PL across the periplasm to preserve PL homeostasis of the Gram-negative OM (Malinverni and Silhavy 2009, Sutterlin et al. 2016, Thong et al. 2016, Ekiert et al. 2017, Ercan et al. 2018, Hughes et al. 2019, Kamischke et al. 2019).

Malinverni & Silhavy (2009) first proposed the Mla pathway as a retrograde transport system involved in the removal of PLs from the outer leaflet of the OM back to the IM through a series of protein complexes (Malinverni and Silhavy 2009); the lipoprotein MlaA at the OM, now identified as being associated with OmpF/C (Chong et al. 2015, Abellon-Ruiz et al. 2017), a periplasmically located soluble protein, MlaC (Ekiert et al. 2017, Hughes et al. 2019), and an inner membrane (IM) ATP binding cassette (ABC) transporter, MlaFEDB of stoichiometry 2:2:6:2 for MlaF, MlaE, MlaD and MlaB (Ekiert et al. 2017). From analysis of Mla knockouts, Malinverni & Silhavy showed an increase in hepta-acylated LPS within the outer leaflet. This was believed to correspond with increased levels of PL present reported through the activity of the palmitoyl transferase PagP, known to synthesise hepta-acylated LPS using the sn-1 palmitoyl of outer leaflet PLs. This phenotype was shown to be recoverable with upregulation of the OM phospholipase protein PldA, known to dimerise and hydrolyze the sn-1 and sn-2 lipid tails of PLs in the presence of excess outer leaflet PLs. These experiments therefore suggested that Mla knockouts resulted in accumulation of outer leaflet PLs and determined that the Mla system had a role in their removal (Malinverni and Silhavy 2009). Further support for retrograde transport was given by Roier *et al*. (2016) who showed increased levels of IM vesicle (OMV) formation in Mla knockouts, suggesting that the accumulation of PL in the outer leaflet led to OMV formation (Roier et al. 2016). However more recently Kamischke *et al.* (2019) provided evidence to the contrary by showing that *Δmla* mutants have a decreased abundance of OM PLs and accumulate newly synthesised PL at the IM through direct quantification of membrane PLs (Kamischke et al. 2019). This was backed up by Ercan *et al.* (2019) who showed the movement of PLs from MlaD to MlaC (Ercan et al. 2018) whilst our laboratory, using membrane reconstituted MlaFEDB, observed unidirectional transport of PL to MlaC and the upregulation of ATPase activity only in the presence of substrate free MlaC (MlaC-apo) (Hughes et al. 2019), supporting a role not in retrograde transport but rather anterograde transport, the movement of PL towards the OM.

Although the key question regarding directionality has not yet been resolved, other questions also remain, in particular what is the role of ATP hydrolysis in the function of MlaFEDB complex? Previously we showed that MlaFEDB driven transport of PLs from the IM to MlaC in the periplasm occurs independently of ATP hydrolysis (Hughes et al. 2019). Despite this, as an ABC transporter, MlaFEDB functions as an ATPase (Thong et al. 2016). This discrepancy in function is difficult to reconcile, especially considering the large amounts of energy required to remove PLs from a bilayer (Marrink et al. 1998). Establishing a link between the ability of MlaFEDB to hydrolyse ATP and transfer PLs therefore represents an important area of research, especially given the potential role MlaFEDB plays in OM biogenesis.

In this study we therefore sought to address this and understand what processes are driven by ATP hydrolysis. Here we show that MlaFEDB has ATP driven flippase activity for all of the major PLs present within *E.coli*. We show flippase activity functions to translocate PLs from the outer to the inner leaflet of the IM and that in the absence of ATP MlaC accepts PL only from the inner leaflet through MlaFEDB. We propose that this flippase activity acts to maintain homeostasis within the IM by replenishing PLs removed from the IM by MlaC, further supporting the role of MlaFEDB in anterograde PL transport.

## Results

### Reconstitution of MlaFEDB in to fluorescently labelled phospholipid proteoliposomes

The purification of N-terminally hexa-histidine tagged MlaFEDB in dodecylmaltoside (DDM) detergent, with the tag present on the N-terminus of MlaF has been described in detail previously (Hughes et al. 2019). This was incorporated in to liposomes containing *E.coli* polar lipids doped with the fluorophore nitrobenzoxadiazole on the headgroup of phosphatidylethanolamine (MlaFEDB:NBD-PE) via size exclusion chromatography (SEC) (Supplementary Fig. 1a). The mixing of NBD-PE with *E.coli* polar lipids prior to rehydration ensured NBD-PE was incorporated into both leaflets of the liposome bilayer. A single uniform peak was observed which was seen to homogeneously consist of MlaFEDB proteoliposomes which were found to be stable and consist of vesicular structures of ~190 nm as measured by Dynamic light scattering (Supplementary Fig. 1b). Further use of SEC allowed us to generate NBD-PE labelled *E.coli* polar lipid liposomes devoid of protein, here termed PL:NBD-PE, and as such could be used as control samples to compare differences in NBD-PE leaflet distribution. NBD-PE was chosen initially, as PE is the predominant PL in *E.coli*, making up ~70% of the total PL pool.

### MlaFEDB preferentially orientates within liposomes

In order to begin to probe the role of ATP in MlaFEDB function, it was important to elucidate the relative orientations of MlaFEDB within the proteoliposome, e.g. either MlaF or MlaD outward facing. To ascertain this we used proteinase K treatment in combination with western blotting against the hexa-histidine tag at the N-terminus of MlaF to ascertain the ratio of outward MlaF facing to inward facing complex (Fig. 1a). It was found that approximately 90% of the complex preferentially oriented itself with the MlaF ATP binding domain outward facing (accessible to ATP), with ~10 % orientating such that MlaD faces outward (accessible to MlaC). However, as both populations existed, unidirectional study could be achieved by either the addition of ATP or the addition of MlaC to the outside of the proteoliposome (Fig. 1b).

**Figure 1.**
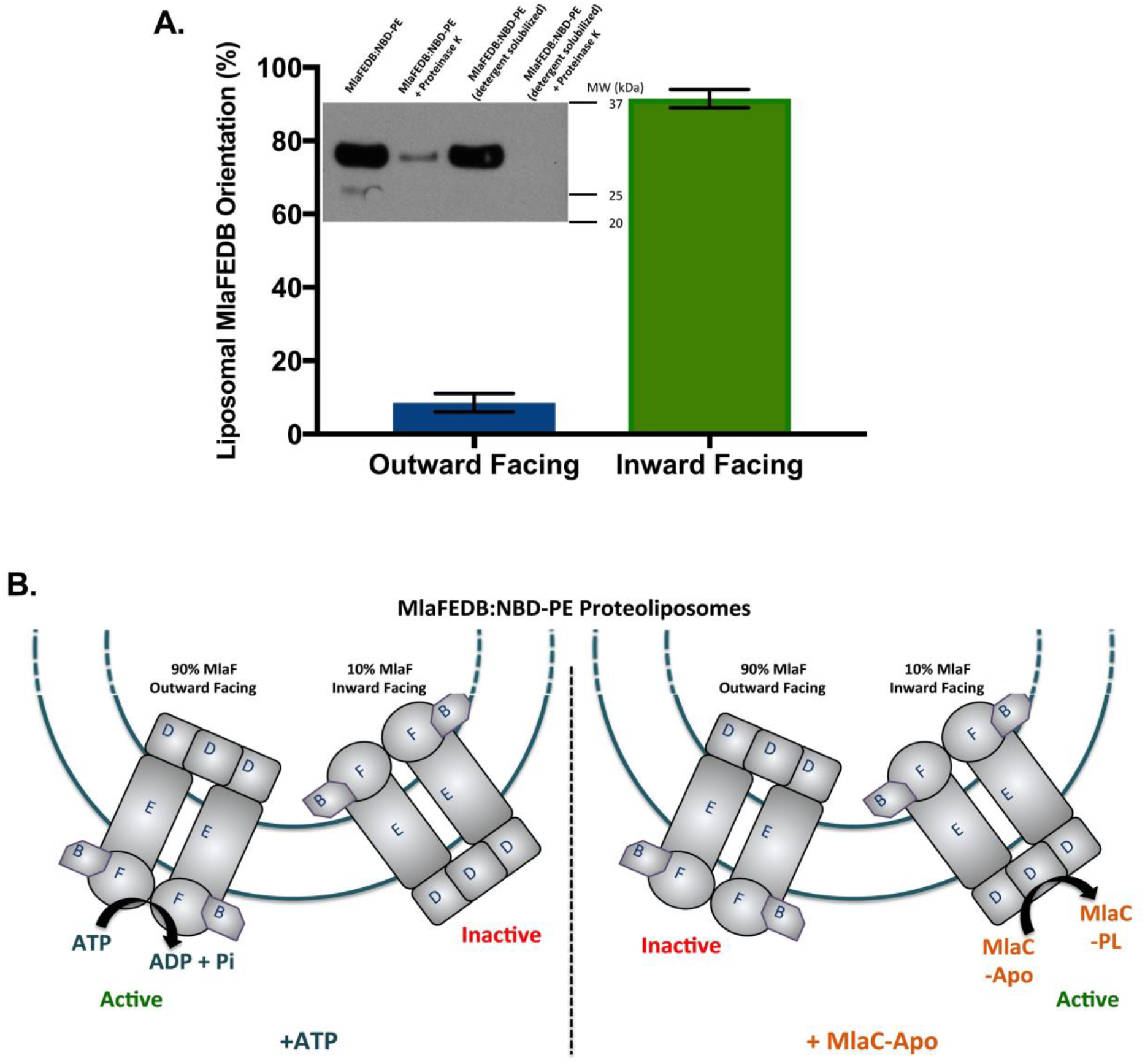
MlaFEDB orientates differentially in liposomes. **A)** MlaFEDB:NBD-PE was treated with proteinase K and western blotting performed, staining with an anti-His antibody, allowing for identification of MlaF. Owing to the impermeability of the MlaFEDB:NBD-PE to proteinase K the orientation of the MlaFEDB complex within the liposome could then be assessed, with any observed anti-His staining following proteinase K treatment resulting from inward facing MlaF. Western blots were then analysed through densitometry to allow quantification of protein orientation. **B)** Schematic diagram showing the preferential orientation of MlaFEDB within liposomes.

### MlaFEDB mediates the transbilayer movement of phospholipids

It is well established that another Gram-negative IM ABC transporter, MsbA, functions as a flippase and is involved in Lipid A transport (Eckford and Sharom 2010). As a functionally similar IM complex, we hypothesised that MlaFEDB may also display the ability to translocate lipids between leaflets of the IM to maintain IM integrity following PL removal. To investigate this potential flippase activity we used a modified lipid flippase assay based on the work of Eckford and Sharom (Eckford and Sharom 2005) to quantify changes in the distribution of fluorescently labelled PLs across proteoliposomal bilayers in response to different treatments. It has already been well established that the Mla pathway shows specificity for the PL tail rather than head group (Thong et al. 2016, Ekiert et al. 2017, Hughes et al. 2019), we hypothesised that the presence of a fluorescent tag would therefore minimally affect activity. Briefly, total fluorescence could be measured at an emission wavelength of 536 nm, with readings equating to the levels of NBD-PE in both the inner and outer leaflets of the membrane. Through the use of the membrane impermeable, fluorescence-quenching agent, sodium dithionite, it is then possible to directly measure inner leaflet fluorescence emitted from NBD-PE alone. Final treatment with Triton X-100 solubilises the proteoliposome, promoting quenching of the inner leaflet and allowing a determination of background fluorescence. As such, it is possible to directly measure the levels of NBD-PE within both the inner and outer leaflets of the membrane.

To ascertain if MlaFEDB performs flippase activity, MlaFEDB:NBD-PE was incubated at 25°C in the presence or absence of ATP for 1 hr. As ATP is unable to cross the liposomal bilayer and was added after proteoliposome formation, any change in distribution would result from the population of MlaFEDB with the MlaF component outward facing only (Fig. 1b). Fluorescence changes observed across the membrane were then measured. In the absence of ATP, NBD-PE was seen to be evenly distributed across the inner and outer leaflets of the MlaFEDB:NBD-PE proteoliposome, with ~48 % NBD-PE present in the inner leaflet following dithionite quenching (Fig. 2a), suggesting that in the absence of ATP no flippase activity was observed, consistent with proteoliposome PL distribution observed in other systems (Eckford and Sharom 2010). However, in the presence of ATP we observed a significant shift in this distribution, with the percentage of inner leaflet NBD-PE decreasing to ~20 % (Fig. 2a). This large change in fluorescence distribution is highly indicative of a MlaFEDB driven ATP dependent translocation of NBD-PE from the inner to the outer leaflet of the proteoliposome.

**Figure 2.**
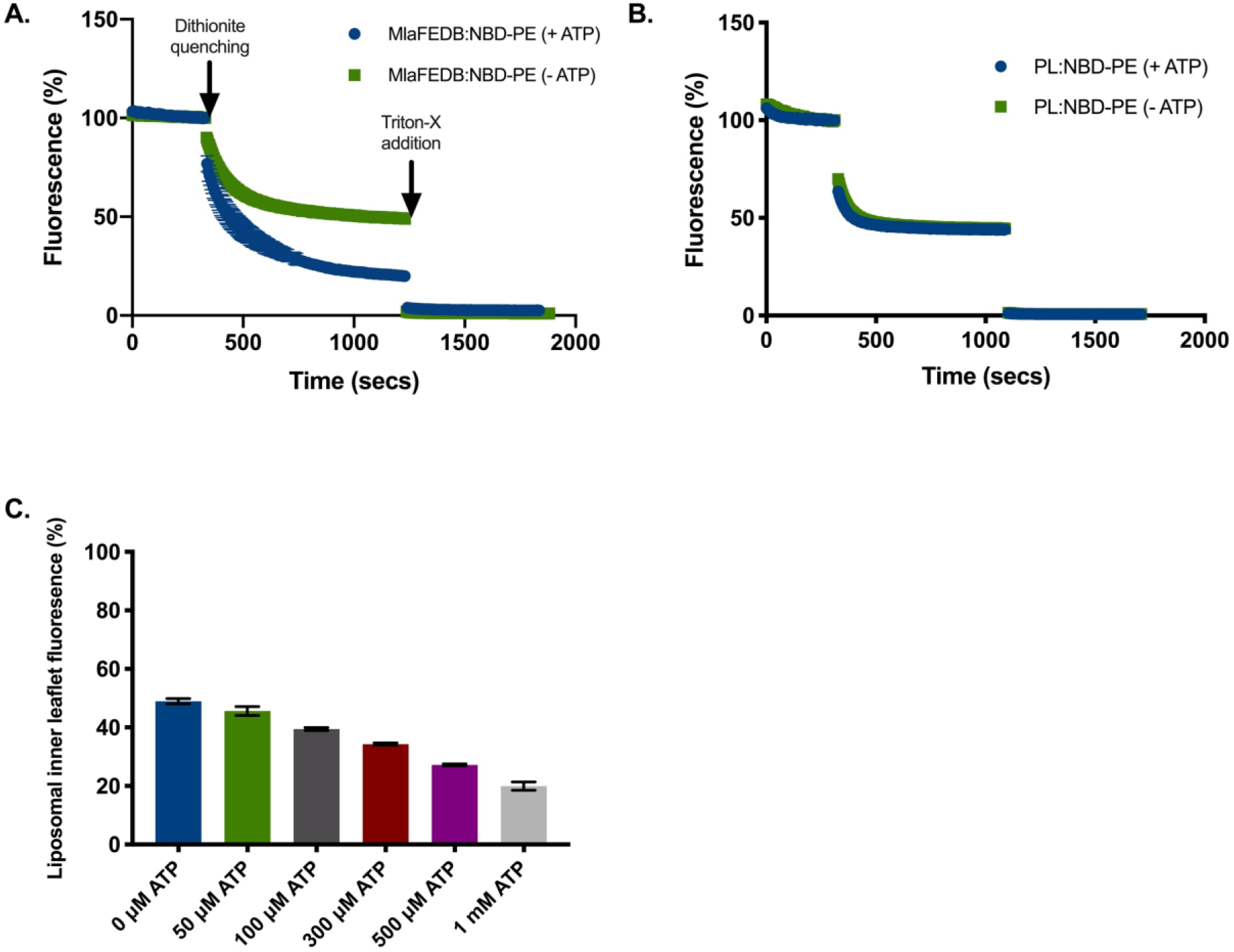
MlaFEDB has phospholipid flippase activity. **A)** MlaFEDB:NBD-PE proteoliposomes were incubated in the presence or absence of 1 mM ATP and a regenerating system for 1 hour. NBD-PE fluorescence was then monitored. Proteoliposomes were then incubated with 4 mM Sodium dithionite (first interruption in trace), with subsequent fluorescence levels observed being representative of inner leaflet NBD-PE levels. Background fluorescence levels were then established through the addition of 1% (w/v) Triton X-100. In the presence of ATP levels of inner leaflet NBD-PE were substantially reduced as compared to untreated proteoliposomes, indicating flippase activity. **B)** The flippase assay as described above was performed using PL:NBD-PE liposomes. ATP did not influence the distribution of NBD-PE across the leaflets of the liposome. **C)** MlaFEDB:NBD-PE flippase assays were performed in the presence of a range of ATP concentrations from 0 - 1 mM.

To ensure that any observed PL translocation is subject to flippase mediated activity and not due to the intrinsic flip-flop of PLs, PL:NBD-PE was also incubated at 25°C in the presence or absence of ATP for 1 hr. Irrespective of the presence of ATP PL:NBD-PE liposomes were seen to display similar levels of NBD-PE distributed across the inner and outer leaflets of the liposome, with ~47 % NBD-PE present in the inner leaflet following dithionite quenching (Fig. 2b). These levels are comparable to those observed in untreated MlaFEDB:NBD-PE proteoliposomes.

To further assess the ATP dependence of MlaFEDB driven NBD-PE translocation we performed translocation assays in the presence of increasing amounts of ATP (0 - 1 mM), with the flippase activity measured over the course of an hour. Translocation of NBD-PE from the inner to the outer leaflet of the proteoliposome was seen to occur in an ATP concentration dependent manner, with increasing levels of ATP resulting in increased movement of NBD-PE to the outer proteoliposomal leaflet (Fig. 2c & Supplementary Fig. 2).

### MlaFEDB flippase activity is dependent upon ATP hydrolysis

Through the use of non-hydrolysable ATP analogues, GTP, and ATPase null mutants we were able to further examine the ATP-dependence of the MlaFEDB driven flippase activity.

To test whether flippase activity is stimulated by the binding of ATP or by the hydrolysis of ATP we analysed the flippase activity of MlaFEDB:NBD-PE proteoliposomes in the presence of ATPγS, a non-hydrolysable analogue of ATP. Previously we have shown MlaFEDB was still able to retain PL transport function, transferring PLs to MlaC in the presence of ATPγS (Hughes et al. 2019). Here in the presence of ATPγS we saw no significant change in the percentage of fluorescent PL (47%) within the inner leaflet, a level comparable to the quantity of fluorescent PL observed in the absence of ATP (47%), suggesting that flippase activity does not occur. Furthermore, this lack of PL flipping was mimicked in the presence of 1 mM GTP, with levels of proteoliposomal inner leaflet fluorescence remaining at around 47 % in both the presence and absence of GTP (Fig. 3a & Supplementary Fig. 3).

**Figure 3.**
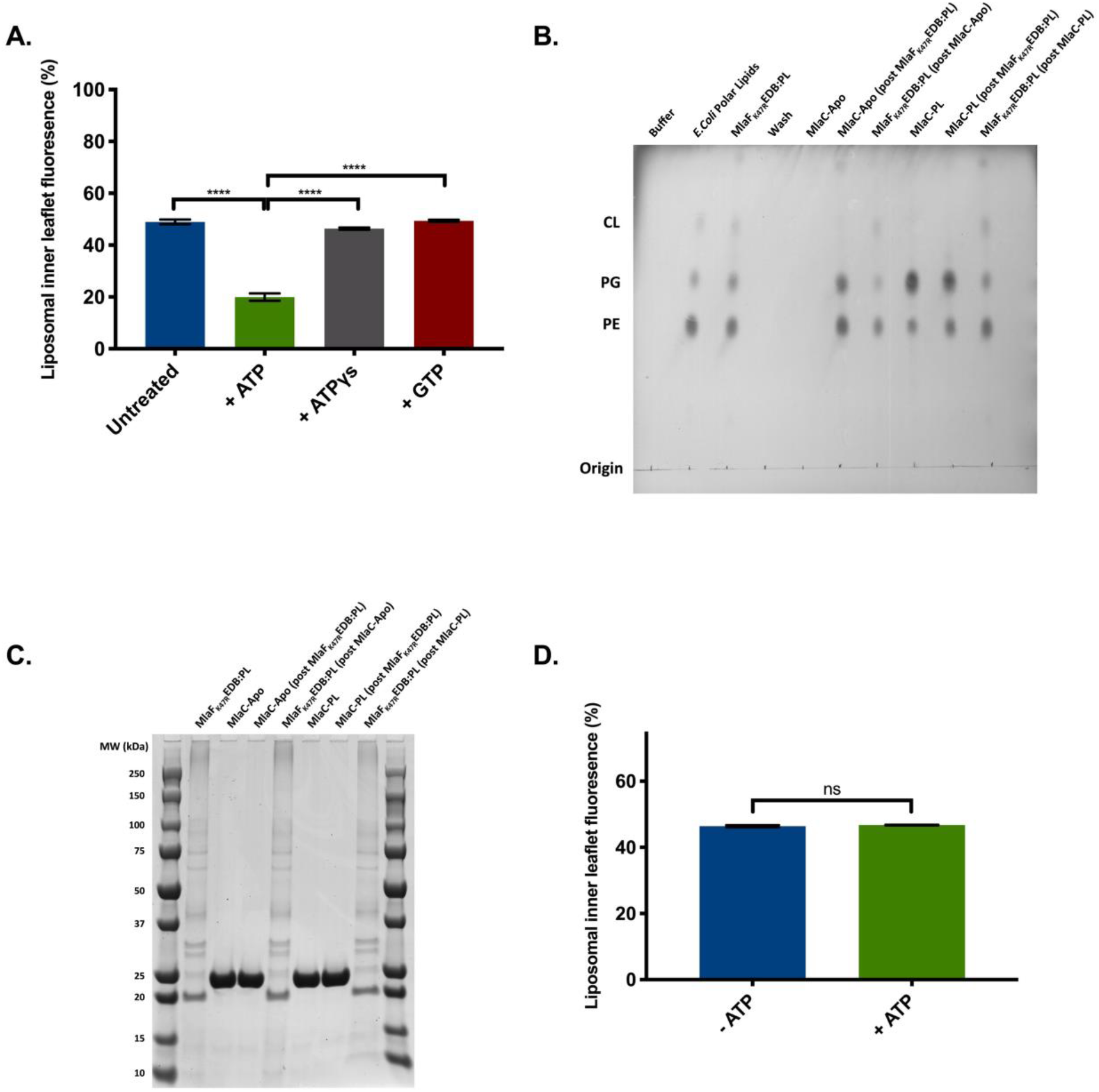
MlaFEDB flippase activity is dependent upon ATP hydrolysis. **A)** MlaFEDB:NBD-PE proteoliposomes were incubated for an hour in the presence of 1 mM ATP, 1 mM ATPγS, 1 mM GTP, or left untreated. Following incubation, the levels of NBD-PE distributed across the leaflets of the proteoliposome were determined through dithionite quenching, allowing us to assess flippase activity. Bars represent the average results from 3 independent flippase assays, with standard deviation shown. ****P < 0.0001 as determined through unpaired t-tests. **B)** Thin-layer chromatogram showing the movement of PLs between MlaF_K47R_EDB and MlaC. **C)** SDS–PAGE of the samples shown in **B** confirming separation of the various species following incubation. **D)** MlaF_K47R_EDB reconstituted into *E.coli* polar lipid liposomes containing 0.3 % (w/w) NBD-PE were incubated either in the presence or absence of ATP and monitored for flippase activity through the use of dithionite quenching. Levels of inner leaflet NBD-PE were quantified for 3 individual assays, error bars represent standard deviation. Ns = no significance.

To further probe the functional independence of MlaFEDB driven PL transport and flippase activity we utilised the MlaFEDB ATPase null mutant, MlaF_K47R_EDB, a non-functional MlaF walker A motif mutation (Thong et al. 2016). Here we purified MlaF_K47R_EDB to homogeneity (Supplementary Fig. 4a) and again showed its lack of inherent ATPase activity (Supplementary Fig. 4b). Through binding to metal chelate resin, we were able to stably reconstitute MlaF_K47R_EDB into *E.coli* polar lipid liposomes, thus allowing us to probe lipid transport through incubation with MlaC-apo and analysis via thin layer chromatography, as we have shown previously for MlaFEDB (Hughes et al. 2019). Despite its lack of ATPase activity MlaF_K47R_EDB retained the ability to transfer PLs to MlaC-apo, whilst no obvious loss of PL from MlaC-PL to MlaF_K47R_EDB was noted (Fig. 3b,c), consistent with our previous observations for MlaFEDB (Hughes et al. 2019). Loading of MlaC-apo with PLs directly from the liposome was discounted as we have previously shown this does not occur (Hughes et al. 2019). However, regardless of its ability to transport PLs to MlaC, MlaF_K47R_EDB lacked flippase activity (Fig. 3d & Supplementary Fig. 5). This is in line with its ATPase inactivity; with the levels of inner leaflet NBD-PE remaining consistent (47%) in both the presence and absence of ATP.

Together these results demonstrate the necessity of ATP hydrolysis for MlaFEDB driven flippase activity and highlight how MlaFEDB mediates two distinct functions, PL transfer to MlaC and lipid translocation between the leaflets of the IM.

### MlaFEDB is able to flip all major species of E.coli phospholipid

To address whether MlaFEDB shows flippase preference for specific PL species present within the bacterial IM, we probed MlaFEDB flippase activity in the presence of headgroup NBD labelled phosphatidylglycerol (MlaFEDB:NBD-PG) and TopFluor® labelled cardiolipin (MlaFEDB:TopFluor®-CL), NBD headgroup labelled cardiolipin was not available. MlaFEDB displayed the ability to flip both phosphatidylglycerol and cardiolipin from the inner to the outer leaflet of the proteoliposome (Fig. 4 & Supplementary Fig. 6a,b). Whilst the directionality of flippase activity remained constant for NBD-PE, NBD-PG, and TopFluor-CL, the levels at which flippase activity occurred differed between PL species. MlaFEDB was able to translocate NBD-PE and NBD-PG at similar rates in the presence of 1 mM ATP, with levels of inner leaflet fluorescence decreasing to ~18 % for both species. However, the extent to which cardiolipin was translocated between leaflets by MlaFEDB was significantly reduced, with around 35 % of Topfluor®-CL remaining in the inner leaflet following ATP incubation over the same time period (Fig. 4). Until now it has remained unclear whether the Mla pathway plays a role in cardiolipin transport, with Thong et al suggesting MlaC is unable to bind it (Thong et al. 2016), whilst we previously provided evidence for binding (Hughes et al. 2019). With the current observation of CL leaflet flipping by MlaFEDB, albeit at a lower level to PG and PE, it more firmly suggests the Mla pathway is able to transport CL. Whether the decreased levels of CL flipping are due to a preference for the diacyl chain containing PE and PG rather than the teta-acyl CL remains unclear however due to having to use an alternate CL fluorophore which could impact on rate. The results however demonstrate that MlaFEDB is able to function as a flippase for all three major PL species found within the bacterial membrane. Furthermore, the observation of flipping from the inner leaflet to the outer leaflet when presented with ATP on the exterior of the MlaFEDB proteoliposome corresponds to a bulk movement of PL towards the MlaF protein and hence within the cell envelope the movement of PLs from the outer to the inner leaflet of the IM.

**Figure 4.**
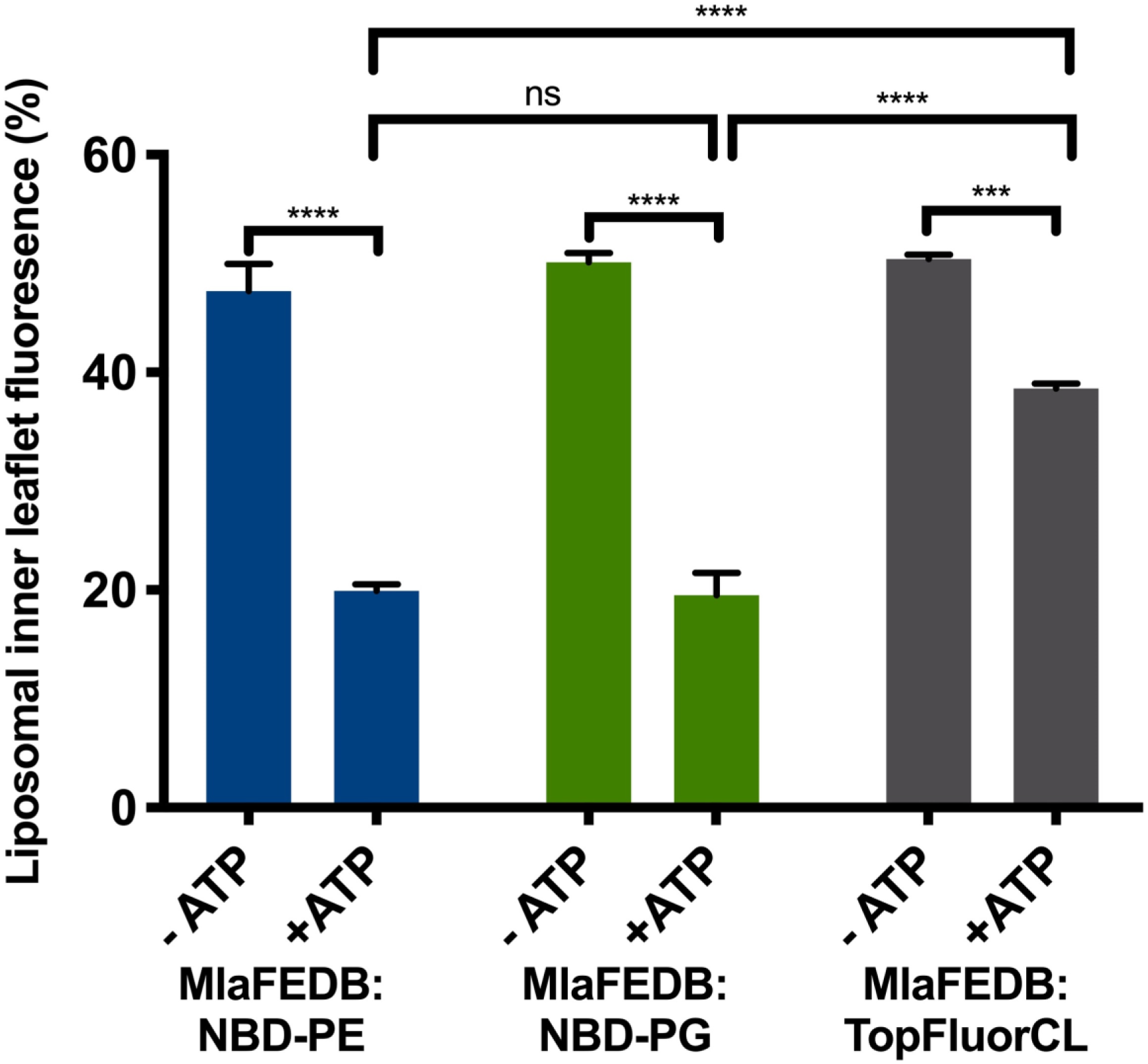
MlaFEDB mediated flippase activity occurs in the presence of all major *E.coli* phospholipids. *E.coli* polar lipid proteoliposomes containing reconstituted MlaFEDB and either 0.3 % (w/w) NBD-PE, 0.3 % (w/w) NBD-PG, or 0.3 % (w/w) TopFluorCL were incubated for an hour in the presence or absence of 1 mM ATP and a regenerating system. The levels of inner leaflet fluorescent lipid was determined through dithionite quenching. All datasets represent the average results from 3 independent flippase assays, with standard deviation shown. Ns: no significance, **P < 0.01, ***P < 0.001, ****P < 0.0001 as determined through unpaired t-tests.

### MlaFEDB flippase activity correlates with MlaC phospholipid loading and transport towards the outer membrane

Following our observation that MlaFEDB mediates the flipping of PLs in a unidirectional fashion from the outer to the inner leaflet of the IM, a process which appears counterintuitive to our previous study showing MlaFEDB functions to extract PL from the IM and donates it to MlaC-apo and infers a shuttling of PLs towards the OM (Hughes et al. 2019), we sought to understand this process in more detail.

With the knowledge that ~10% of the MlaFEDB complexes within the proteoliposome orient with the MlaD component outward facing (Fig. 1a,b), incubation of MlaFEDB:NBD-PE proteoliposomes with MlaC-apo yielded information regarding the leaflet from which PLs are exported to MlaC (Fig. 5a & Supplementary Fig. 7). Upon incubation of MlaC-apo with MlaFEDB:NBD-PE proteoliposomes lacking ATP, a significant reduction in the inner leaflet level of NBD-PE was observed compared to the outer leaflet. As these assays were performed in the absence of ATP it can be inferred that this change in ratio was due to transfer to MlaC rather than flippase activity, certainly no flippase activity was previously noted in the absence of ATP. In contrast, the addition of MlaC-PL to MlaFEDB:NBD-PE fluorescently labelled proteoliposomes was seen to have no obvious effect on the levels of inner leaflet fluorescence. This being in line with the levels observed in untreated MlaFEDB:NBD-PE proteoliposomes. These results therefore indicate that MlaC affects the levels of NBD-PE within the inner leaflet. MlaC mediated removal of PLs from the outer leaflet could not be measured using this method as the fluorescence signal for the outer leaflet was inferred from the total fluorescence minus inner leaflet fluorescence and therefore comprised of both MlaC-NBD-PE bound and outer leaflet NBD-PE.

**Figure 5.**
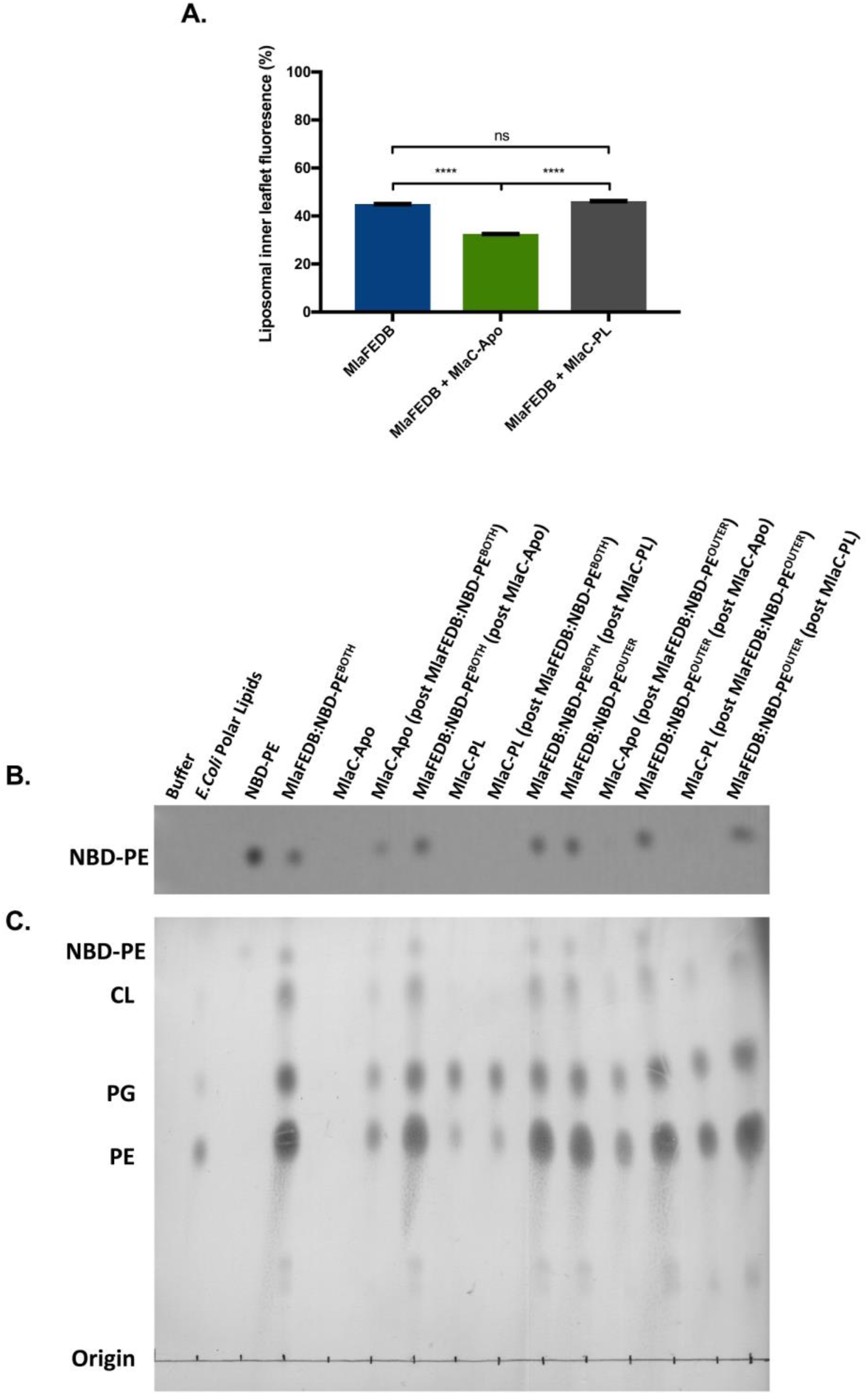
MlaFEDB flippase activity facilitates PL transfer onto MlaC. **A)** MlaFEDB:NBD-PE proteoliposomes were incubated for an hour at a 1:1 molar ratio with either MlaC-Apo, MlaC-PL, or left untreated. The transbilayer distribution of NBD-PE following this incubation was determined through the use of dithionite quenching. Only in the presence of MlaC-Apo was inner leaflet fluorescence decreased. Bars represent the average results from 3 independent flippase assays, with standard deviation shown. Ns: no significance, ****P < 0.0001 as determined through unpaired t-tests. **B)** To further assess the ability of MlaC to accept PLs from the inner leaflet of MlaFEDB proteoliposomes we reconstituted MlaFEDB into *E.coli* polar lipid liposomes in which only the outer leaflet contained NBD-PE at a final concentration of 0.3 % (w/w) (MlaFEDB:NBD-PE^OUTER^). Either MlaFEDB:NBD-PE^OUTER^ or MlaFEDB reconstituted into polar lipid liposomes containing 0.3 % (w/w) NBD-PE distributed across both leaflets (MlaFEDB:NBD-PE^BOTH^) were incubated with either MlaC-Apo or MlaC-PL. The transfer of NBD-PE between these species was then assessed through fluorescence visualization after separation via TLC. Labels indicate what protein lipids have been extracted from, following incubations with other proteins denoted by brackets. **C)** Identical TLC as shown in **b** following staining with 10 % (w/v) phosphomolybdic acid (PMA) labelling mimics that of **b**.

To further investigate which bilayer leaflet PLs were extracted from we introduced NBD-PE into the outer leaflet of MlaFEDB *E.coli* polar lipid proteoliposomes (MlaFEDB:NBD-PE^OUTER^) using the approach of Chiantia *et al*. (Chiantia et al. 2012) and compared PL transfer capabilities to that of MlaFEDB proteoliposomes in which both leaflets were labelled (MlaFEDB:NBD-PE^BOTH^) (Supplementary Fig. 8). We could then incubate these proteoliposomes with MlaC-apo and through the use of TLC and fluorescence imaging monitor the transfer of NBD-PE (Fig. 5b). Following incubation with MlaFEDB:NBD-PE^BOTH^, as expected MlaC-apo became loaded with NBD-PE by the clear fluorescence observed on the TLC plate, thus indicating transfer of NBD-PE between MlaFEDB and MlaC. However, on incubation of MlaC-apo with MlaFEDB:NBD-PE^OUTER^ almost no fluorescence was observed. Density mapping estimated the signal to be approximately 10% of that following incubation with MlaFEDB:NBD-PE^BOTH^. PL transfer was still seen to occur as staining of the TLC plate with PMA highlighted MlaC-apo was loaded with PLs (Fig. 5c). This therefore suggests uptake occurred from the inner liposomal leaflet only. We attribute the ~10% signal to the observation that IM labelling with NBD-PE wasn’t 100%. We observed ~10% being incorporated into the inner leaflet thus some NBD-PE could gain access to MlaC. Incubation of MlaC-PL with MlaFEDB:NBD^OUTER^ or MlaFEDB:NBD^BOTH^ resulted in no change in observed fluorescence and an inability to donate or receive any PLs, suggesting the binding of PL to MlaC is sufficiently tight that exchange cannot occur.

These results therefore show that in the presence of MlaC-apo and in the absence of ATP, PLs are extracted from the inner leaflet of the MlaFEDB proteoliposome. In this scenario, where MlaC can only interact with the population of MlaFEDB with the MlaD component outward facing suggests that within the context of the cell that, in the absence of ATP, MlaC is loaded with PLs solely from the inner leaflet of the IM.

## Discussion

The mechanism of Mla driven PL transport remains a controversial subject, with conflicting views within the literature regarding the directionality of PL movement (Powers and Trent 2019). Recently we showed that in isolation MlaFEDB transfers PLs from MlaFEDB to MlaC and suggested that the Mla system might mediate the flux of PLs towards the OM (Hughes et al. 2019). However, the role of ATP within this process remained unclear, as we observed PL transport occurring without the need for an energy source. Despite this, whole cell studies performed by Kamishke and colleagues provided evidence to suggest that ATP is needed for Mla mediated transport of PLs toward the OM (Kamischke et al. 2019). As such, elucidating how ATP influences PL transport represents one of the key questions remaining regarding Mla mediated activity. Here we have attempted to reconcile these discrepancies and provide evidence to suggest MlaFEDB functions as a flippase in addition to its PL transport activity, and is able to flip PLs from the outer to the inner leaflet of the IM. We also show that on interaction with MlaFEDB and in the absence of ATP, MlaC gets loaded with PLs solely from the inner leaflet of the IM.

The significant MlaFEDB flippase activity observed in this study initially seemed at odds with our previous findings in which we observed an ATP independent export of PLs into the periplasm via MlaC (Hughes et al. 2019). Especially confusing was the observation of PL flipping towards the MlaF component, which in the context of the cell represents the movement of PLs from the outer to the inner leaflet, the opposite direction to PL export. These results in isolation provide credence to the idea that MlaFEDB could function in a retrograde transport fashion by redistributing PLs across both bilayers following influx from the OM. However, our previous observation showing PL export to MlaC (Hughes et al. 2019), the recently published structures of MlaFEDB adopting an exporter fold (Coudray et al. 2020, Mann et al. 2020), and our current findings casts doubt on this model. But how then can flippase activity and export be rationalised?

Work from our group, in addition to studies performed by Ekiert *et al.* (2017), have outlined the PL binding mechanism of MlaC (Ekiert et al. 2017, Hughes et al. 2019), with fatty acid chains buried deep within the MlaC binding pocket, whilst the head group protrudes from the protein surface. PLs must therefore be orientated on MlaD such that when MlaC binds the acyl tails are presented to it. Our observation that MlaC is loaded with inner leaflet PLs in the absence of ATP can therefore be rationalised as the PLs in this leaflet are already positioned in the correct orientation with their tails presented (Fig. 6) and thus represents both an energetically and spatially favourable mechanism of PL transfer. In this scenario, PLs could be directly exported to MlaC via a continuous flow driven by the affinity of MlaC for PLs and its prevalence. Further support for this model is given by the recently published structure of the *A. Baumannii* MlaFEDB complex in which a channel appears to be observed through MlaE from PLs bound to MlaFEDB on the inner leaflet side directly to MlaD (Mann et al. 2020). What then is the role of the ATPase driven flippase action? This at first glance is somewhat more difficult to reconcile. Is this just an aberrant effect due to the use of fluorescently labelled PLs? Or if real, why is there a need for PLs to be flipped when they can be extracted directly from the inner leaflet without ATP hydrolysis? We propose it is real and is backed up by the recently released structure of *E. coli* MlaFEDB (Coudray et al. 2020). This complex was solved in the apo configuration with PLs observed within the core of the MlaE dimer. Although presented as the “outward-open” configuration, neither PL was observed in the tail up configuration as would be expected for export to MlaD and more likely represents the MLaFEDB complex prior to flipping of PLs to the inner leaflet. What then is the role of the flippase? We propose it is there to enable PLs to be extracted from both leaflets, thus maintaining membrane homeostasis. If extraction only occurred from the inner leaflet, curvature would ensue (McMahon and Gallop 2005, Zimmerberg and Kozlov 2006, Devaux et al. 2008). Flippase activity would therefore enable transfer between bilayers and would thus allow extraction from both leaflets, the assumption being that two PLs are exported from the inner leaflet for every one flipped from the outer leaflet, resulting in a net loss of one PL from each leaflet. Within the cell ATP levels are sufficiently high that this process would likely be continual but presumably the export and flippase activity of MlaFEDB must be coordinated. However, in our *in vitro* study the observation of flippase activity and export apparently occurring independently seems somewhat at odds with this, and in truth we currently can’t provide a clear answer for this observation but it must be noted we have previously shown that ATPase activity is upregulated in the presence of MlaC-apo implying coordination between the two processes does occur. It must also be noted that MlaB is a STAS domain containing protein. STAS domains have previously been implicated in regulation (Sharma et al. 2011), therefore it is feasible to suggest that the STAS domain containing MlaB might also participate in this. Indeed, Thong *et al*. (2016) have previously outlined a role for MlaB in influencing MlaFEDB ATPase activity (Thong et al. 2016), a notion recently reinforced by Kolich et al. (Kolich et al. 2020), however the exact mechanism of this regulation remains to be elucidated. As such it could be that some form of dysregulation by MlaB has occurred in our *in vitro* experiments that led to the disconnect between flippase activity and export. Indeed Mann et al (2020) noted only 50% of their cryo-EM class averages having MlaB bound to both MlaF proteins, the remaining only had one copy (Mann et al. 2020). It is therefore not inconceivable to assume the same could happen here.

**Figure 6.**
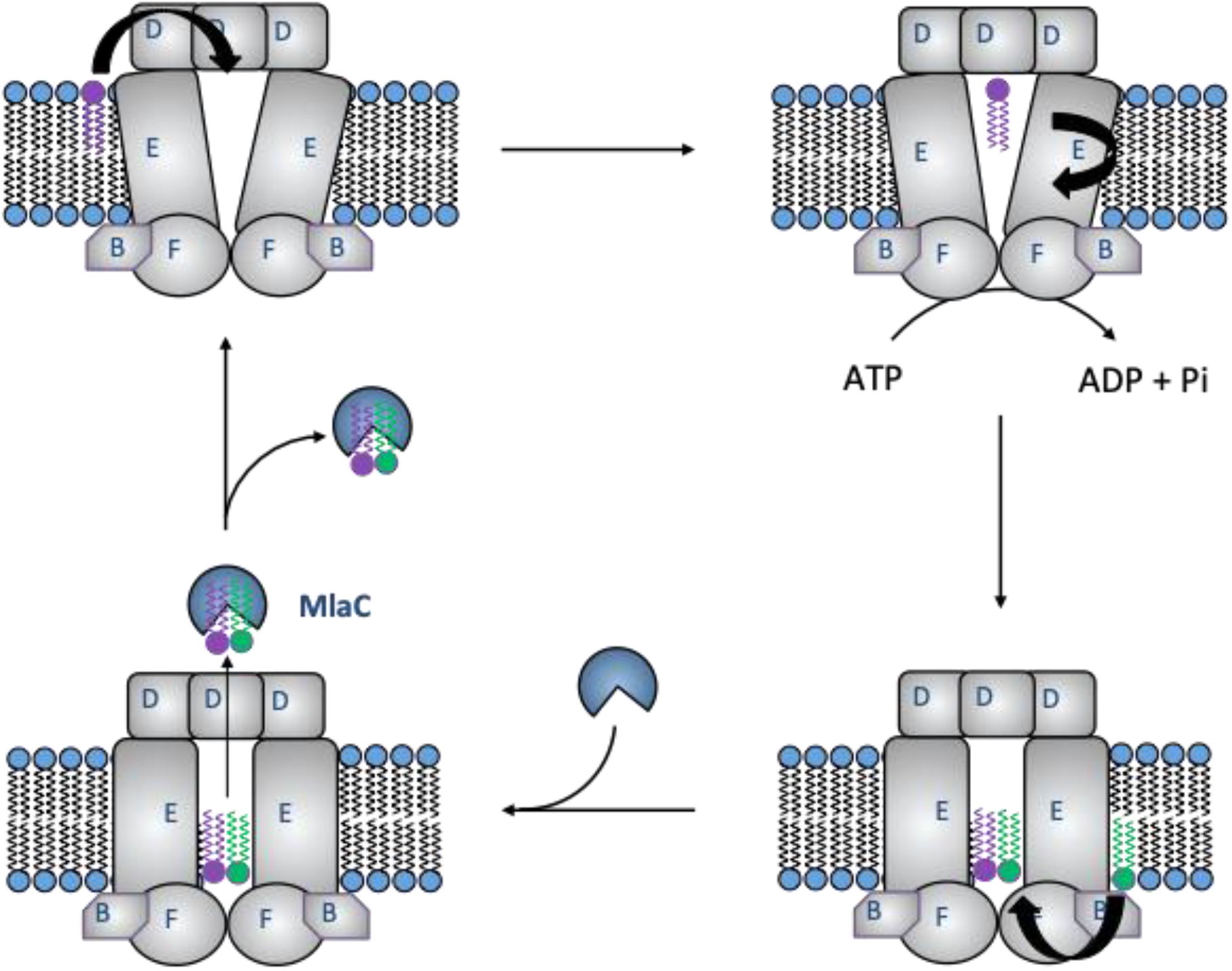
Schematic diagram highlighting how MlaFEDB flippase activity correlates with PL transport onto MlaC. Within the context of the cell, MlaFEDB flippase activity mediates the translocation of PLs from the outer to the inner leaflet of the IM. We suggest that this flippase activity is important in maintaining the integrity of the IM. We suggest that MlaC is loaded solely with inner leaflet PLs via MlaD, as such MlaFEDB flippase activity is required to replenish the lipids in the inner leaflet that are lost upon transfer to MlaC. In light of new structural evidence regarding MlaFEDB, we suggest flippase activity plays an essential role in orientating PLs to allow for the effective transfer onto MlaC. ATP hydrolysis may allow for outer leaflet PLs present within the complex to translocate its inner leaflet side, the subsequent change in conformation may then also allow for an extra inner leaflet PL to access the MlaE cavity. This would create a situation in which two PLs are poised for movement through the MlaE central cavity and loading onto MlaD. This PL movement through the cavity may rely on the action of MlaC stripping PLs previously bound to MlaD, which in turn increases MlaFEDB ATP hydrolysis, thus facilitating increased PL loading onto MlaD.

In summary, this study provides evidence to suggest that MlaFEDB exhibits PL flippase activity, and can facilitate the transbilayer movement of phosphatidylethanolamine, phosphatidylglycerol and cardiolipin from the outer to the inner leaflet of the IM. We also show that in the absence of ATP MlaFEDB mediates the transfer of PLs from the inner leaflet of the IM to MlaC within the periplasm. These findings highlight MlaFEDB as the first generic PL flippase discovered in Gram-negative bacteria and provide further mechanistic details to support the Mla driven anterograde PL transport process.

## Materials and Methods

### Expression and purification of MlaFEDB

MlaFEDB was expressed and purified following methodology as described by (Ekiert et al. 2017). Briefly, Rosetta 2 (DE3) cells (Novagen) were transformed with the plasmid pBE1196 (complete *mlaFEDCB* operon containing a hexa-histidine tag at the C terminus of MlaF). MlaF_K47R_EDB mutants were generated through site-directed mutagenesis using the MlaFEDB pBE1196 plasmid as target DNA and purified via the same protocol as MlaFEDB. Firstly, overnight cultures were grown to an OD600 of 0.9 with shaking at 37 °C. Addition of arabinose to a final concentration of 0.1% (w/v) with 4 hours shaking incubation at 37 °C was used to induce MlaFEDB expression. The cells were pelleted through centrifugation at 6,000*g* for 10 mins before resuspension in 50 mM Tris pH 8, 150 mM NaCl, 10 mM imidazole. The cells were then lysed through three passes through an Emulsiflex C3 disruptor (Avestin). Lysate was then subjected to two rounds of centrifugation, the first (20,000*g*, 30 min) to remove the cell debris, and the second (100,000*g*, 30 min) to pellet membranes. The membrane pellets were solubilized for 1 h at 4 °C in 50 mM Tris pH 8, 500 mM NaCl, 10 mM imidazole, 25 mM DDM, before insoluble material was removed by centrifugation at 100,000*g* for 30 mins. Solubilized membranes were then bound to a 5-ml Ni-NTA column (HisTrap, GE Healthcare), which was subsequently washed with 50 mM Tris pH 8, 150 mM NaCl, 50 mM imidazole, 0.5 mM DDM, before bound protein was eluted with 50 mM Tris pH 8, 150 mM NaCl, 250 mM imidazole, 0.5 mM DDM. MlaFEDB containing fractions were then subjected to additional purification on a Superdex 200 size-exclusion column (GE Healthcare) equilibrated in 50 mM Tris pH 8, 150 mM NaCl, 0.5 mM DDM.

### Expression and purification of MlaC

A custom plasmid (pBE1203) containing DNA corresponding to MlaC with an N-terminal hexa-histidine tag followed by a TEV protease cleavage site (Ekiert et al. 2017) was transformed into *E.coli* BL21(DE3) cells. Overnight cultures were grown in lysogeny broth at 37 °C to an OD600 = 0.6 whereupon protein expression was induced by the addition of isopropyl-β-d-thiogalactoside to a final concentration of 1 mM, cells were then continually grown overnight at 15 °C. The cells pelleted through centrifugation at 6,000*g* for 10 mins and then resuspended in 50 mM Tris pH 8, 500 mM NaCl, 10 mM imidazole. Cell pellets were lysed by three passes through an Emulsiflex C3 disruptor (Avestin) and then centrifuged (75,000*g*, 45 min) to remove cell debris. Clarified lysate was then flowed over a 5-ml Ni-NTA column (GE Healthcare), before being washed with 50 mM Tris pH 8, 500 mM NaCl, 50 mM imidazole. Finally, bound protein was eluted with 50 mM Tris pH 8, 500 mM NaCl, 500 mM imidazole. MlaC containing fractions were further purified through size exclusion chromatography on a Superdex 75 column (GE Healthcare) equilibrated in 50 mM Tris pH 8, 150 mM NaCl.

### Lipid removal from MlaC

Generating MlaC-Apo was performed as described previously by (Hughes et al. 2019). Briefly, purified MlaC was bound to a Ni-NTA affinity column (GE Healthcare) and washed with 25 ml of 50 mM Tris pH 8, 500 mM NaCl, 10 mM imidazole, 25 mM β-OG followed by a 1 h static incubation period in the wash buffer. This process of washing and incubation was repeated 3 times before being followed by a final wash with 50 ml of 50 mM Tris pH 8, 500 mM NaCl, 10 mM imidazole. MlaC was then eluted using 50 mM Tris pH 8, 500 mM NaCl, 250 mM imidazole and dialysed against 50 mM Tris pH 8, 150 mM NaCl.

### Formation of proteoliposomes

Proteoliposomal formation was based on methodologies performed by (Eckford and Sharom 2010). Briefly, either solely 10 mg of *E.coli* polar lipids (Avanti Polar Lipids), or doped with either 0.3 % (w/w) NBD-PE, 0.3 % (w/w) NBD-PG, or 0.3 % (w/w) TopFluorCL were dried down under a constant stream of nitrogen. Dried lipid samples were partially solubilized in 250 μL in 50 mM Tris pH 8, 150 mM NaCl, 0.5 mM DDM before 1 mg of purified protein in DDM (MlaFEDB, MlaF_K47R_EDB) was added, producing a final lipid:protein ratio of 10:1 (w/w). Samples were then incubated on ice for 30 mins with occasional mixing before being separated on a Superdex 75 gel filtration column equilibrated in 50 mM Tris pH 8, 150 mM NaCl, 5 mM MgCl_2_. Proteoliposomes eluted in the column void volume, were pooled and adjusted to give a final protein concentration of 0.5 mg/ml.

To solely label the outer leaflet of MlaFEDB proteoliposomes with NBD-PE a method outlined in (Chiantia et al. 2012) was adapted. Briefly, NBD-PE lipids were dried down and resuspended in 100 % Ethanol to produce a stock concentration of 20 mg/ml. These lipids were then added to formed MlaFEDB proteoliposomes as outlined above to produce a final NBD-PE concentration of 0.3 % (w/w). Proteoliposomes were subsequently separated from non-inserted free NBD-PE molecules through separation on a Superdex 75 gel filtration column equilibrated in 50 mM Tris pH 8, 150 mM NaCl, 5 mM MgCl_2_.

In those experiments in which proteoliposomes contained differential leaflet labelling, distinction has been made in which proteoliposomes containing NBD-PE in the outer leaflet alone are termed MlaFEDB:NBD-PE^OUTER^ whilst those proteoliposomes labelled with NBD-PE in both inner and outer leaflets are termed MlaFEDB:NBD-PE^BOTH^.

### Lipid translocation / flippase assay

The flippase activity of various MlaFEDB constructs was determined through measuring differences in the ratios of NBD labelled lipids across the different leaflets of the proteoliposome. Flippase assays were performed using the same methodology regardless of which liposomal/proteoliposomal system was used. Firstly, proteoliposomes were subjected to a variety of different conditions, including incubation with a range of ATP concentrations (0 - 1 mM), 1 mM ATPγS, 1 mM GTP, or either 0.5 mg/ml MlaC-Apo or MlaC-PL. Incubation was performed over an hour period at room temperature, and for flippase specific assays in the presence of an ATP regeneration system (0.5 mM Phosphoenolpyruvate (Sigma-Aldrich) and 100 U ml^−1^ pyruvate kinase (Sigma-Aldrich)). Following incubation, the reaction mixture was transferred to a 96-well fluorescence based assay microplate (Thermo Fisher Scientific) (200 μL reaction volume) and fluorescence emission (NBD-Lipids: excitation at 464 nm, emission at 536 nm; TopFluor-CL: excitation at 498 nm, emission at 507 nm) monitored at 25 °C until a stable baseline was established. 4 mM Sodium Dithionite was added to the reaction mixture and fluorescence again monitored until readings stabilized. Finally, 1 % (w/v) Triton X-100 was added, with fluorescence measured to give a background reading.

### Dynamic light scattering

Dynamic light scattering experiments were performed using a DynaPro® Plate Reader III and DYNAMICS software (Wyatt Technology, Haverhill, UK), using the laser wavelength of 825.4 nm with a detector angle of 150°. Each sample (40 μL) was loaded into a 384-well glass bottom SensoPlate™ (Greiner Bio-One, Germany) in triplicate. Each measurement consisted of 10 scans of 5 s, carried out at 25 °C, with the attenuator position and laser power automatically optimized for size determination (nm).

### ATPase assay

An NADH enzyme-linked assay (Norby 1988) that was adapted for a microplate reader (Kiianitsa et al. 2003) was used to determine either the MlaFEDB or MlaF_K47R_EDB rate of ATP hydrolysis. 0.1 μM of protein, 200 mM NADH (Sigma-Aldrich), 20 U ml^−1^ lactic dehydrogenase (Sigma-Aldrich), 100 U ml^−1^ pyruvate kinase (Sigma-Aldrich), 0.5 mM phosphoenolpyruvate (Sigma-Aldrich) and different ATP (Sigma-Aldrich) concentrations were added to the wells of a 96-well plate (Sigma-Aldrich) and made up in assay buffer (50 mM Tris pH 8.0, 150 mM NaCl, 5 mM MgCl_2_, 0.5 mM DDM) to produce a total final assay volume of 75 μl. Using an Anthos Zenyth 340rt (Biochrom) absorbance photometer equipped with ADAP software the change in absorbance at 340 nm was measured. Reactions were followed for 10–60 min with measurements taken every 15s. Using maximal 340 nm absorbance readings (with 0 μM ATP) and minimal 340 nm absorbance readings (with 0 μM NADH) ATP-hydrolysis rates were determined; the concentration of NADH at each time-point was calculated from these readings, allowing for a linear fit of the reduction in NADH absorbance and conversion to ATP hydrolysis.

### Thin layer chromatography

PL movement within the Mla system was monitored through a process of incubation between different MlaFEDB proteoliposomes and MlaC in either its -Apo or -PL forms followed by separation through SEC. Proteoliposomes including MlaFEDB:PL, MlaF_K47R_EDB:NBD-PE, MlaFEDB:NBD-PE^BOTH^, and MlaFEDB:NBD-PE^OUTER^ were adjusted to 2 mg/ml and incubated in solution with either MlaC-Apo or MlaC-PL at 2 mg/ml for 1 hour. Following incubation, proteoliposomes were separated from MlaC through SEC using a Superdex 75 column (GE Healthcare) equilibrated in 50 mM Tris pH 8, 150 mM NaCl. Samples were then collected for lipid extraction.

2 ml of sample was adjusted to 0.5 mg/ml and added to 2 ml methanol and 1 ml chloroform. Samples were then vortexed for 5 min, incubated for 30 min at 50 °C and vortexed again for 5 min. The mixture was centrifuged (2,000*g*, 10 min), and the lower phase was extracted and evaporated. Dried lipids were resuspended in 100 μl chloroform, and 5 μl was loaded onto a Silica TLC plate (Sigma) and run with a 6.5:2.5:1 (chloroform:methanol:acetic acid) solvent. The TLC plate was dried for 30 min, stained with 10% (w/v) phosphomolybdic acid in ethanol and heated until staining occurred. For detection of NBD-lipid fluorescence Silica TLC plates were left unstained and visualised using an Amersham Imager 680 blot and gel imager (GE Healthcare).

### Western blotting

In order to determine the orientation of MlaFEDB once reconstituted into polar lipid liposomes, MlaFEDB:NBD-PE was subjected to Proteinase K digestion. 1 mg of MlaFEDB:NBD-PE was incubated with 0.05 mg/ml Proteinase K (Thermo Fisher) for 10 mins at 4 °C. In order to ensure the stability of the proteoliposomes, a control sample of MlaFEDB:NBD-PE was treated with 1 % (v/v) Triton X-100 before being subjected to Proteinase K digestion as outlined above. Proteinase K digested proteoliposomes in both unsolubilzed and Triton X-100 solubilized conditions were compared to samples that were left undigested. All samples were then boiled at 95 °C for 10 mins at a 1:1 (v/v) ratio in 2x Laemmli loading dye (Sigma) before being electrophoresed through a 4-12% NuPAGE Bis-Tris polyacrylamide gel (BioRad). Protein bands were then transferred from the gels onto 0.2 μm polyvinylidene fluoride (PVDF) membranes (Bio-Rad, CA, USA) using semi-dry electroblotting system (Trans-Blot Turbo Transfer System, Bio-Rad). Membranes were blocked for 1 hour with 5% (w/v) dried milk powder dissolved in 10 mM Tris pH 8, 150 mM NaCl, 0.1% (v/v) Tween-20 (TBST). Membranes were then incubated for 4 hours with anti-His hexahistidine antisera conjugated to horseradish peroxidase (HRP) (Sigma) diluted in TBST at a ratio of 1:5000. Following 3 washes in TBST, membranes were incubated with Rabbit Anti-Mouse IgG secondary antisera (Abcam) diluted at a ratio of 1:25000 in TBST. An enhanced chemiluminescence (ECL) detection kit (Merck Millipore) was used to develop the membranes, with chemiluminescence signals visualized using a cp1000 film processor (Agfa).

## Acknowledgements

We thank D. Ekiert and G. Bhabha for discussion and kindly providing Mla plasmid constructs. This research was supported by BBSRC grants BB/P009840/1 and BB/S017283/1 (GWH, PS, & TJK) and BBSRC PhD funding through the Midlands Integrative Biosciences Training Partnership (PJW, SAN, & BFC).

## Author Contributions

GWH, PS, SAN, PJW, BFC and TJK all participated in the conception and design of the work. GWH, PS, SAN, PJW, BFC and TJK all participated in the data acquisition, analysis or interpretation of the work. GWH, PJW, BFC and TJK were involved in writing and editing the manuscript. All authors approved the final version submitted for publication.

## Figure Legends

**Supplementary Figure 1.**
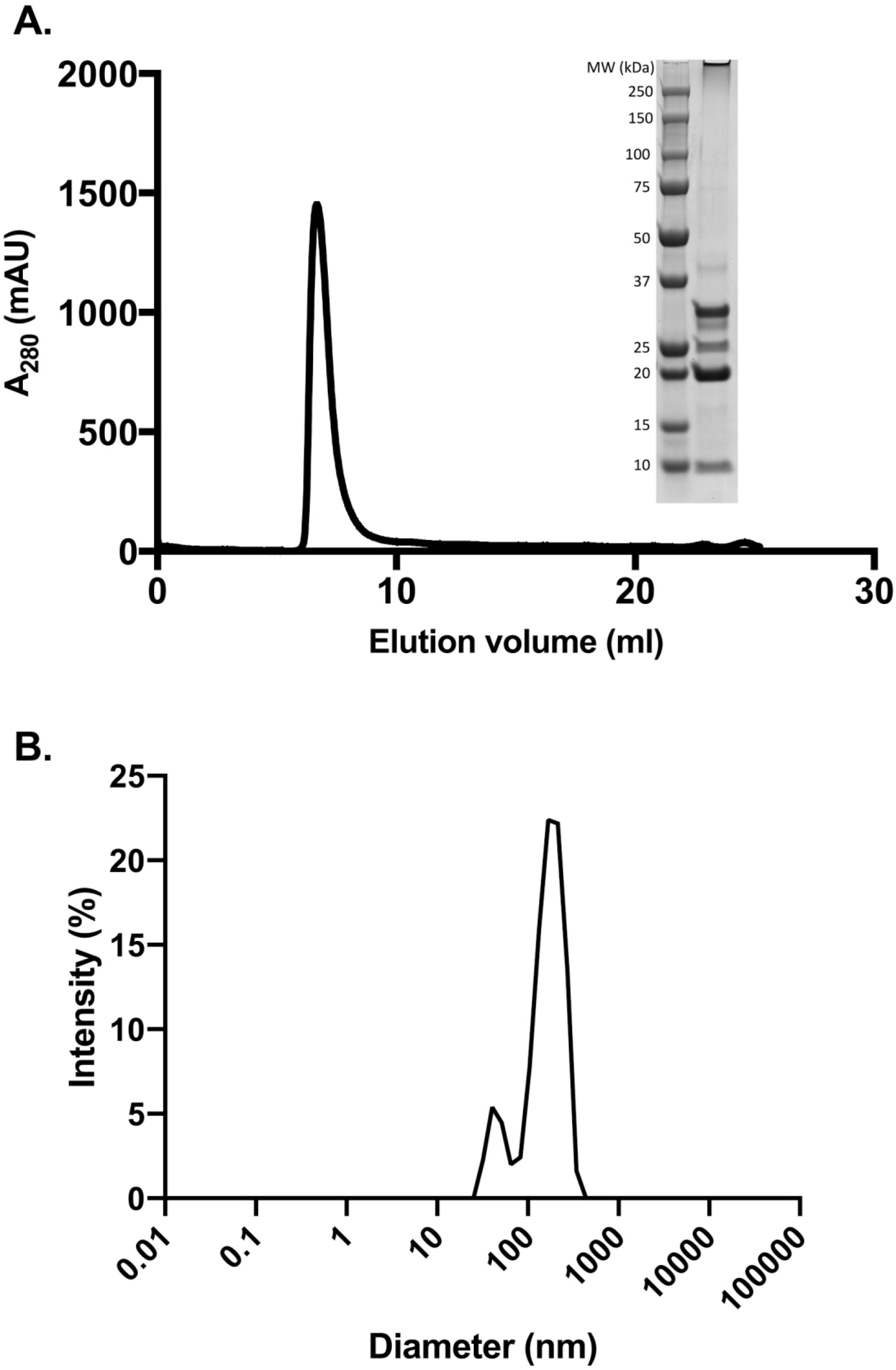
MlaFEDB reconstitution into liposomes. **A)** Size-exclusion chromatogram and SDS-PAGE showing the reconstitution of MlaFEDB into proteoliposomes consisting of *E.coli* polar lipids containing 0.3 % (w/w) NBD-PE (MlaFEDB:NBD-PE). **B)** DLS measurement showing size distribution of MlaFEDB:NBD-PE proteoliposomes.

**Supplementary Figure 2.**
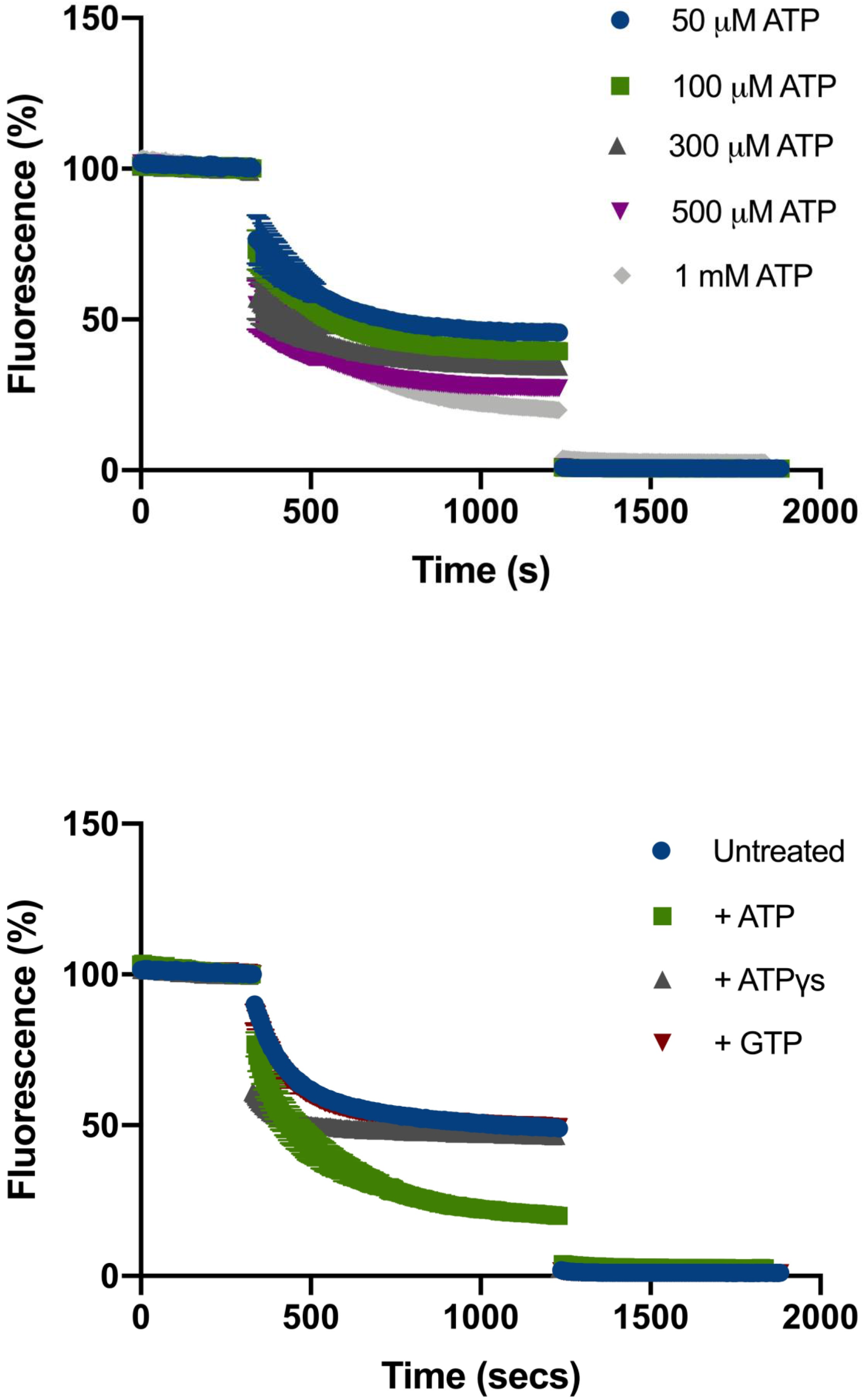
MlaFEDB flippase activity increases in an ATP concentration dependent fashion. Kinetic fluorescence plot displaying the effect of dithionite quenching on the flippase activity assay corresponding to Fig. 2c.

**Supplementary Figure 3.**
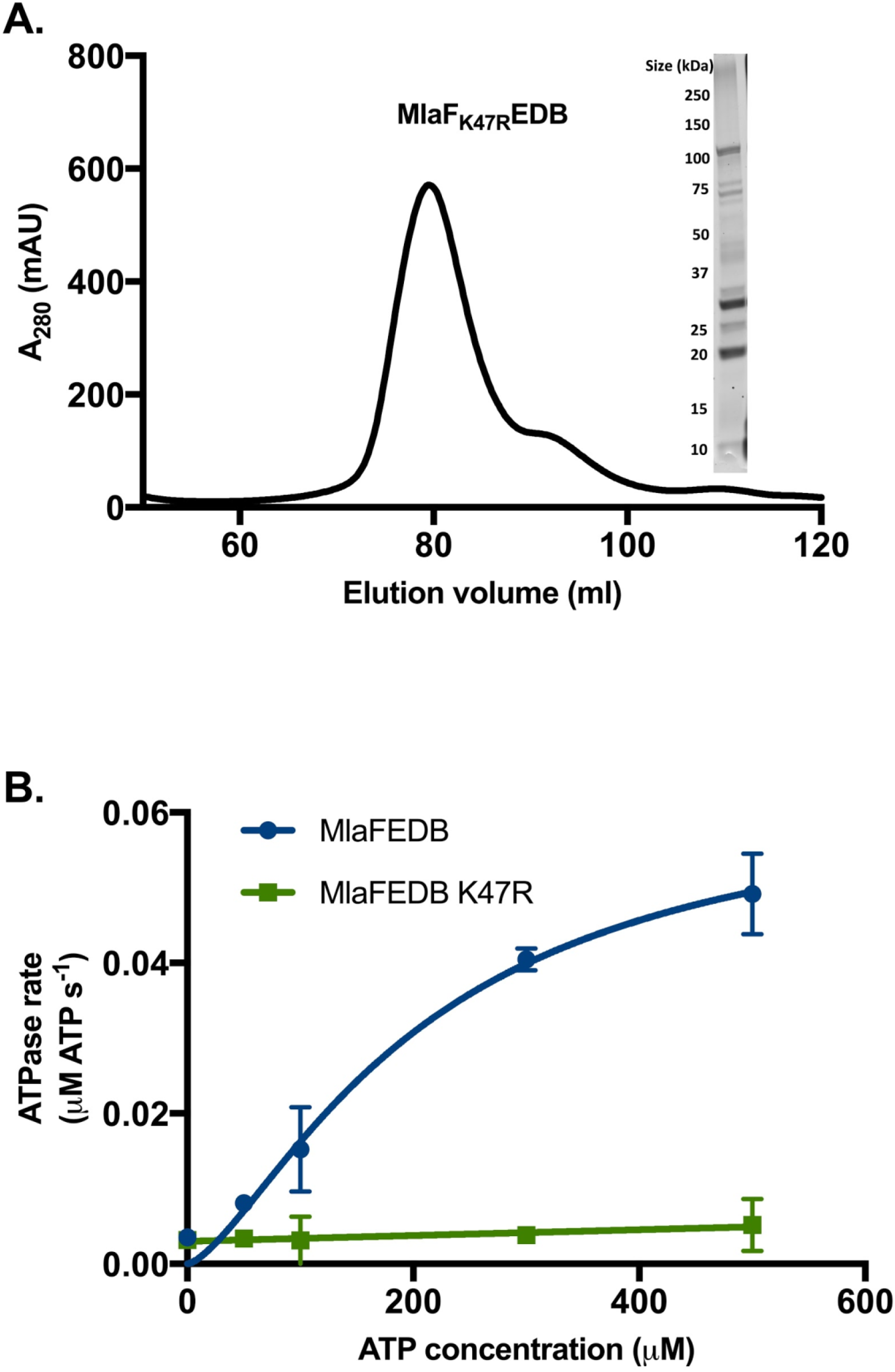
The flippase activity of MlaFEDB relies on ATP hydrolysis. Flippase assay plot corresponding to Fig. 3a displaying full kinetic fluorescence data.

**Supplementary Figure 4.**
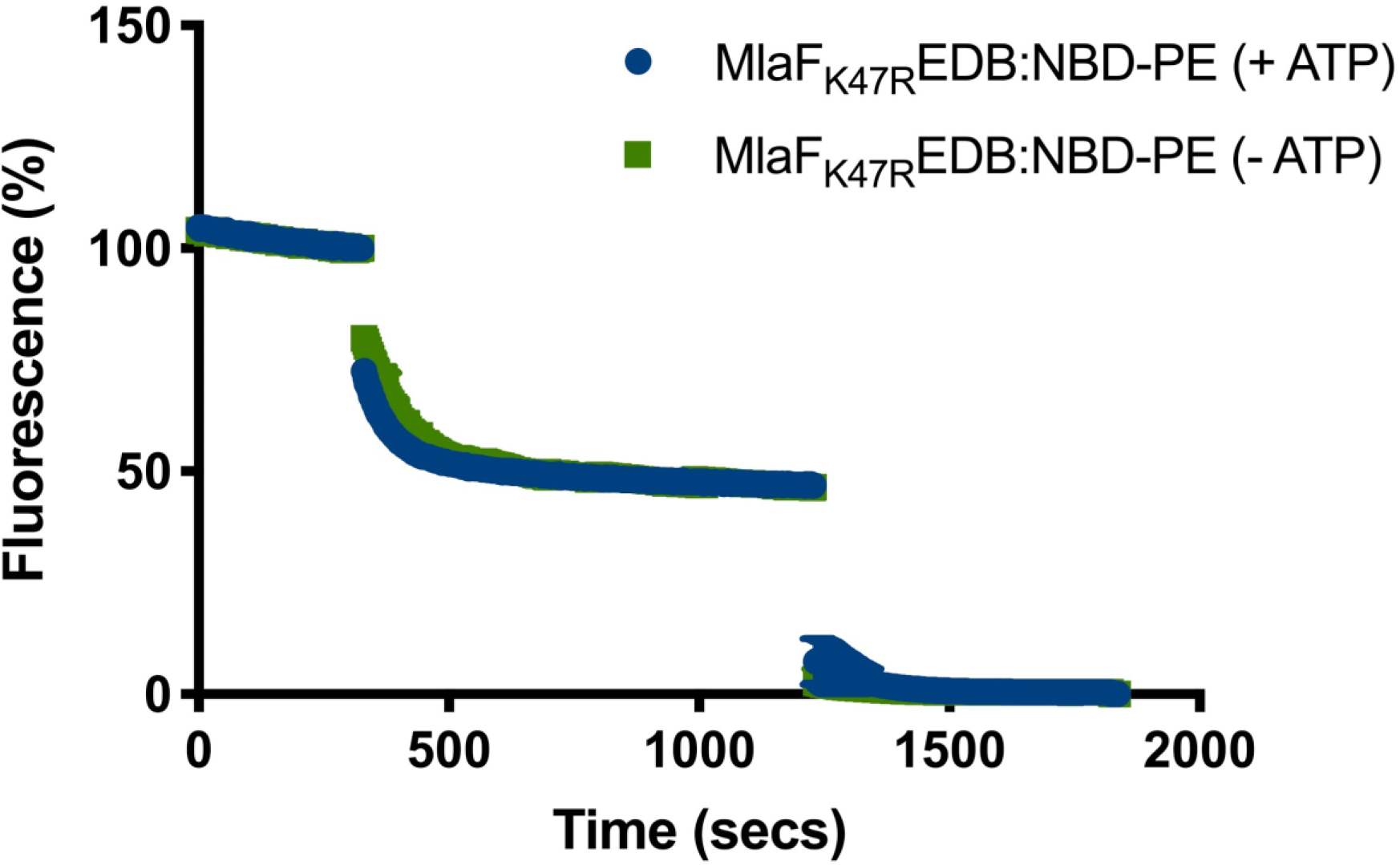
MlaF_K47R_EDB is unable to hydrolyse ATP. **A)** Size-exclusion chromatogram and corresponding SDS-PAGE of detergent-solubilized MlaF_K47R_EDB yielding a single complex. **B)** Enzyme coupled ATPase assay of MlaF_K47R_EDB (0.1 μM) compared to MlaFEDB (0.1 μM) performed in detergent micelles (0.05% DDM). The average ATP-hydrolysis rates were plotted against the ATP concentrations and fitted against an expanded Michaelis–Menten equation that includes a term for the Hill coefficient (n). For MlaFEDB, V_max_ = 0.05 μM ATP s−1, kcat = 0.5 ± 0.1 s−1, K_m_ = 200.0 ± 44.6 μM and n = 1.81 ± 0.35. The error bars indicate the s.d.; n = 3 independent experiments.

**Supplementary Figure 5.**
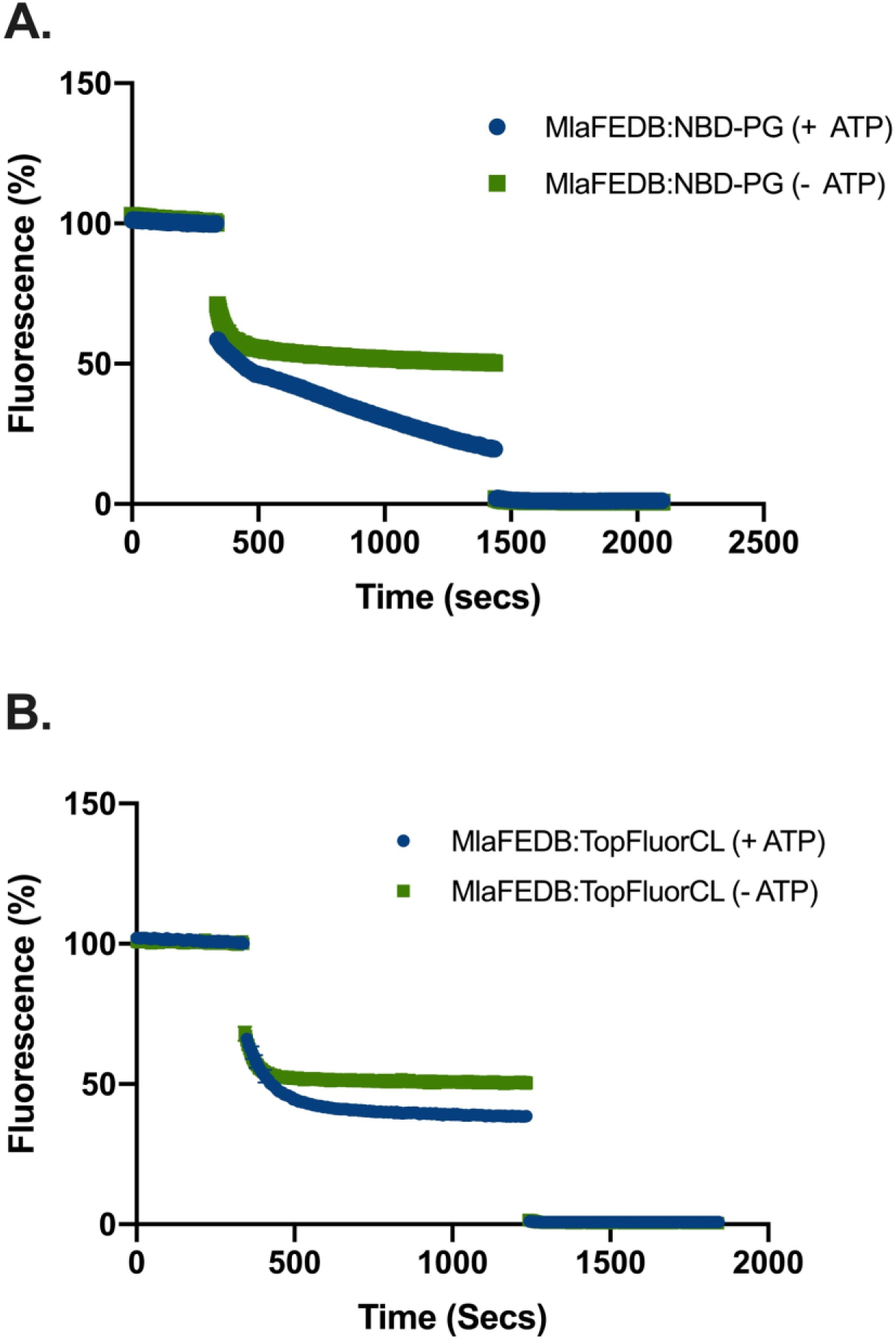
MlaF_K47R_EDB displays no flippase activity. Kinetic fluorescence plot displaying the effect of dithionite quenching on the flippase activity assay corresponding to Fig. 3d.

**Supplementary Figure 6.**
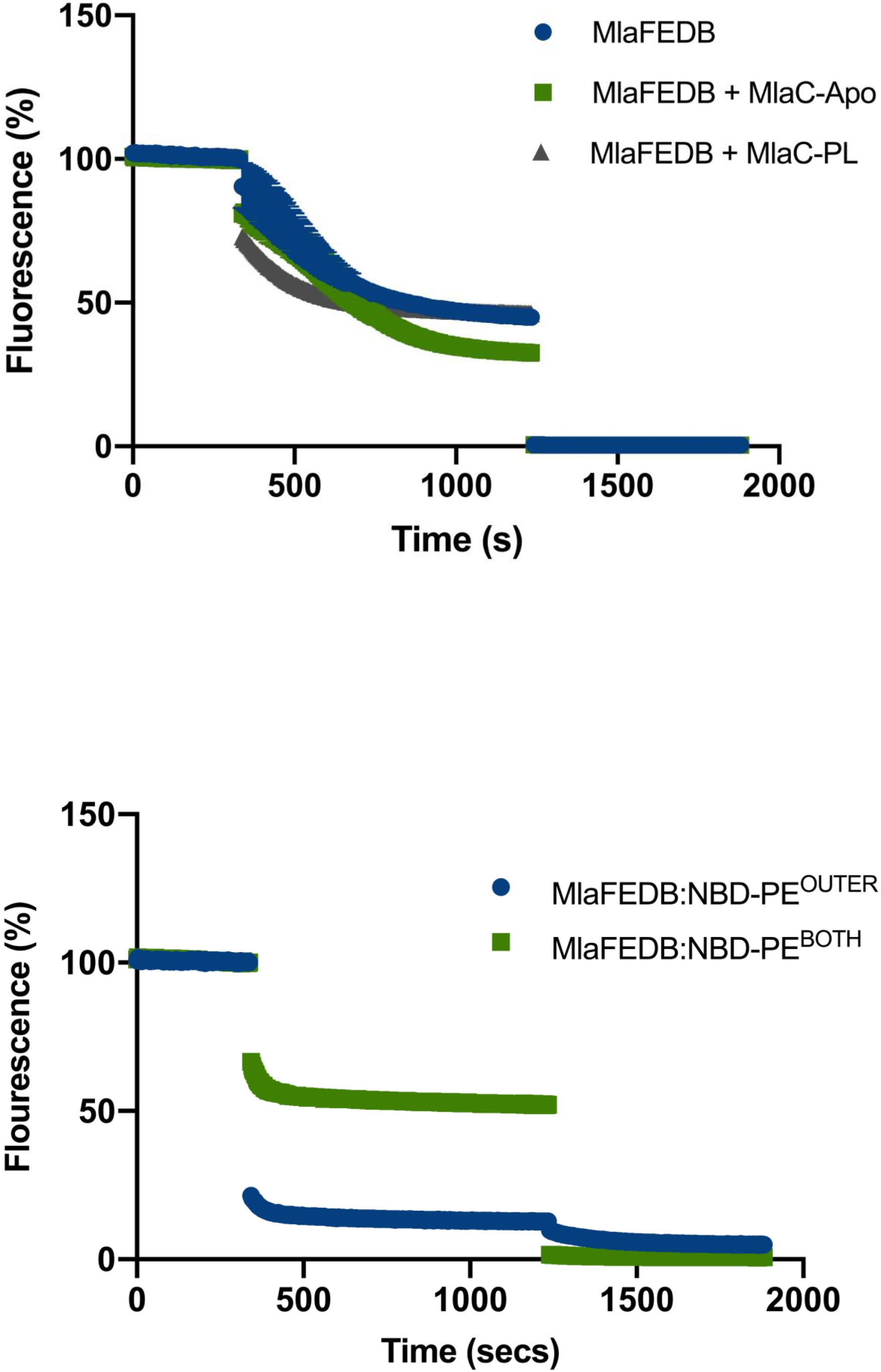
Lipid dependence of MlaFEDB flippase activity. **A)** MlaFEDB:NBD-PG flippase assay corresponding to Fig. 4 showing the effect of dithionite quenching on fluorescence levels. **B)** Kinetic fluorescence plot displaying the effect of dithionite quenching on the MlaFEDB:TopFluorCL flippase activity assay corresponding to Fig. 4.

**Supplementary Figure 7.**
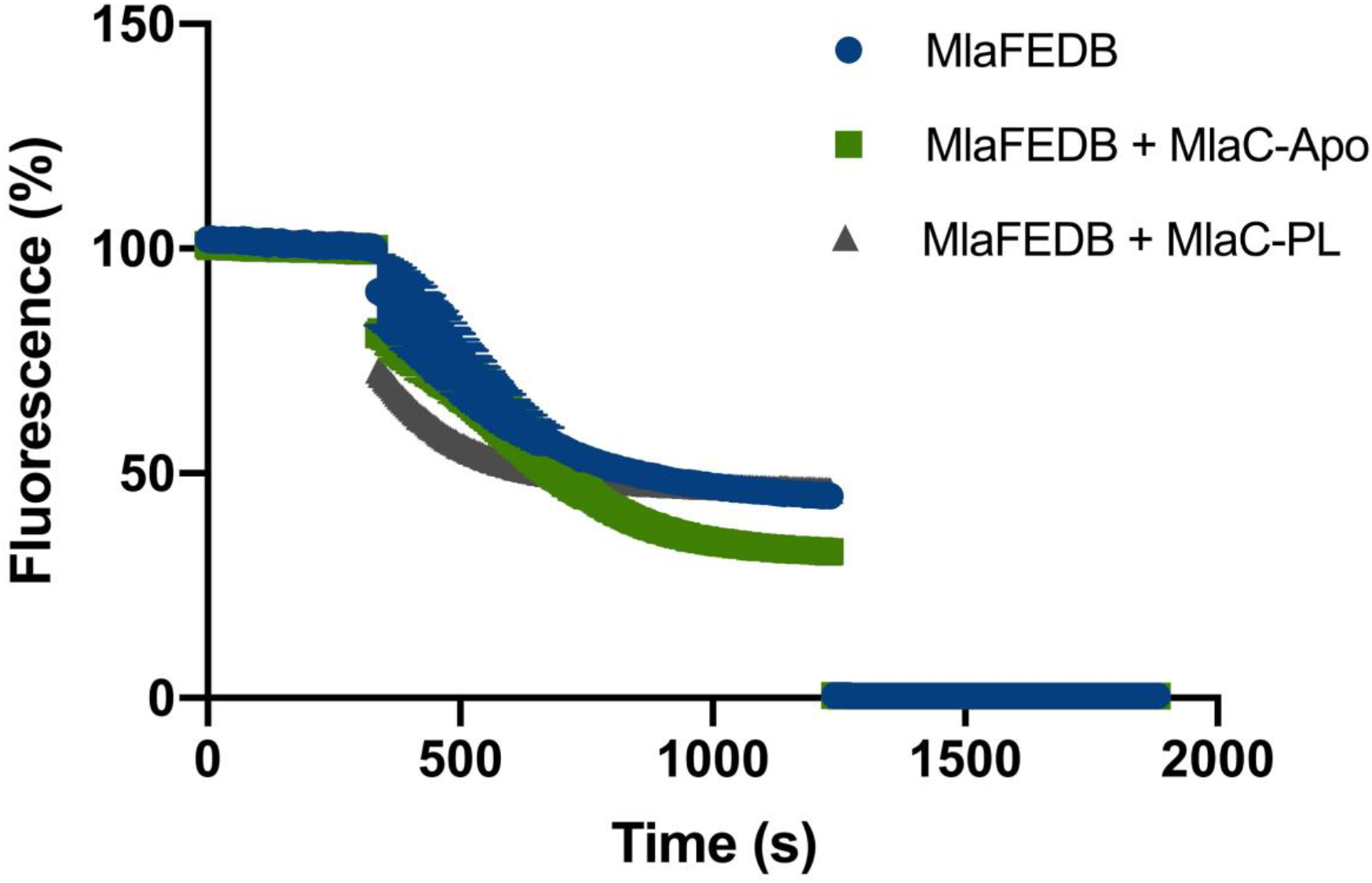
MlaC-Apo affects the transbilayer lipid distribution of MlaFEDB proteoliposomes. Graph corresponding to Fig. 5a showing changes in fluorescence in response to the dithionite quenching of MlaFEDB:NBD-PE proteoliposomes in response to incubation with either MlaC-Apo or MlaC-PL.

**Supplementary Figure 8.**
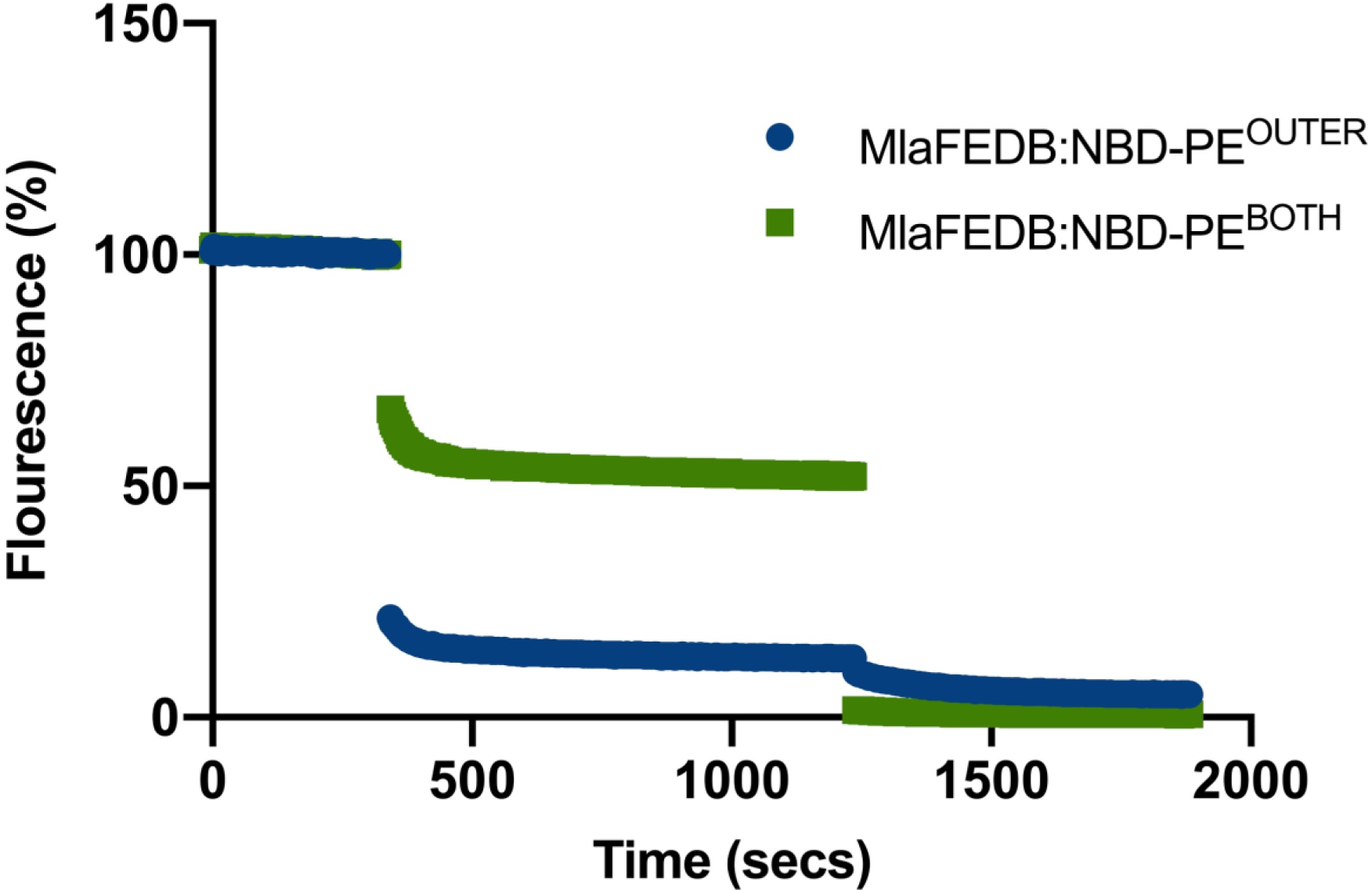
Outer leaflet labelling of MlaFEDB proteoliposomes with NBD-PE. MlaFEDB reconstituted into *E.coli* polar lipid liposomes was incubated with 0.3 % (w/w) NBD-PE in ethanol, resulting in incorporation into the outer leaflet of the proteoliposome only. Through dithionite quenching the levels of NBD-PE in the respective leaflets of these proteoliposomes was assessed and compared to that of MlaFEDB proteoliposomes with NBD-PE distributed across both leaflets. ~ 95 % of NBD-PE in MlaFEDB:NBD-PE^OUTER^ proteoliposomes was found in the outer leaflet. This was compared to MlaFEDB:NBD-PE^BOTH^ proteoliposomes in which NBD-PE was seen to be distributed relatively evenly across both leaflets.

**Supplementary Figure 9.**
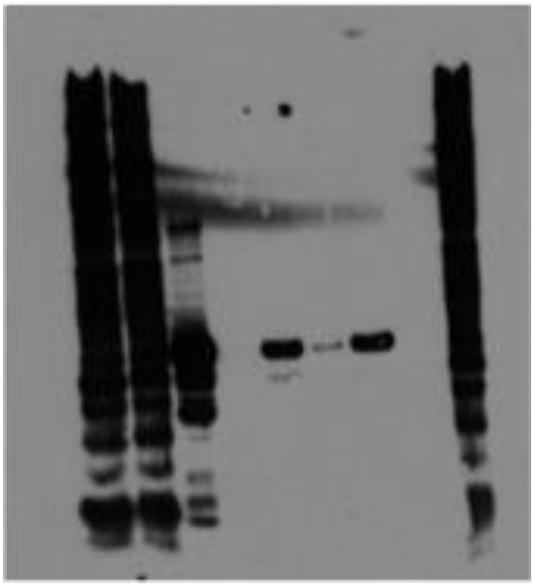
Raw image corresponding to figure 1A.

**Supplementary Figure 10.**
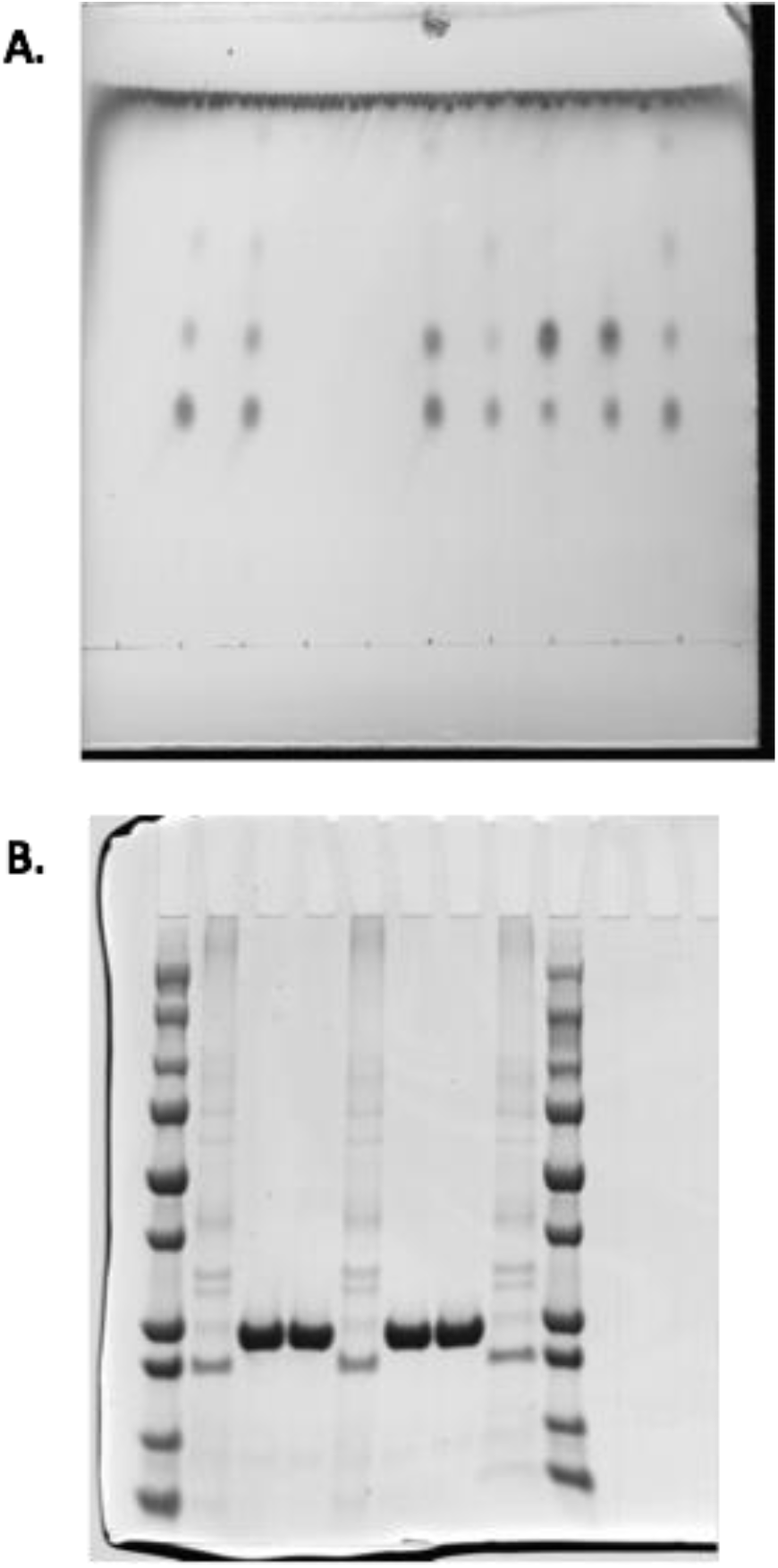
Raw images corresponding to figure 3. **A)** Complete image of TLC plate corresponding to figure 3B. **B)** Raw SDS-PAGE image relating to figure 3C.

**Supplementary Figure 11.**
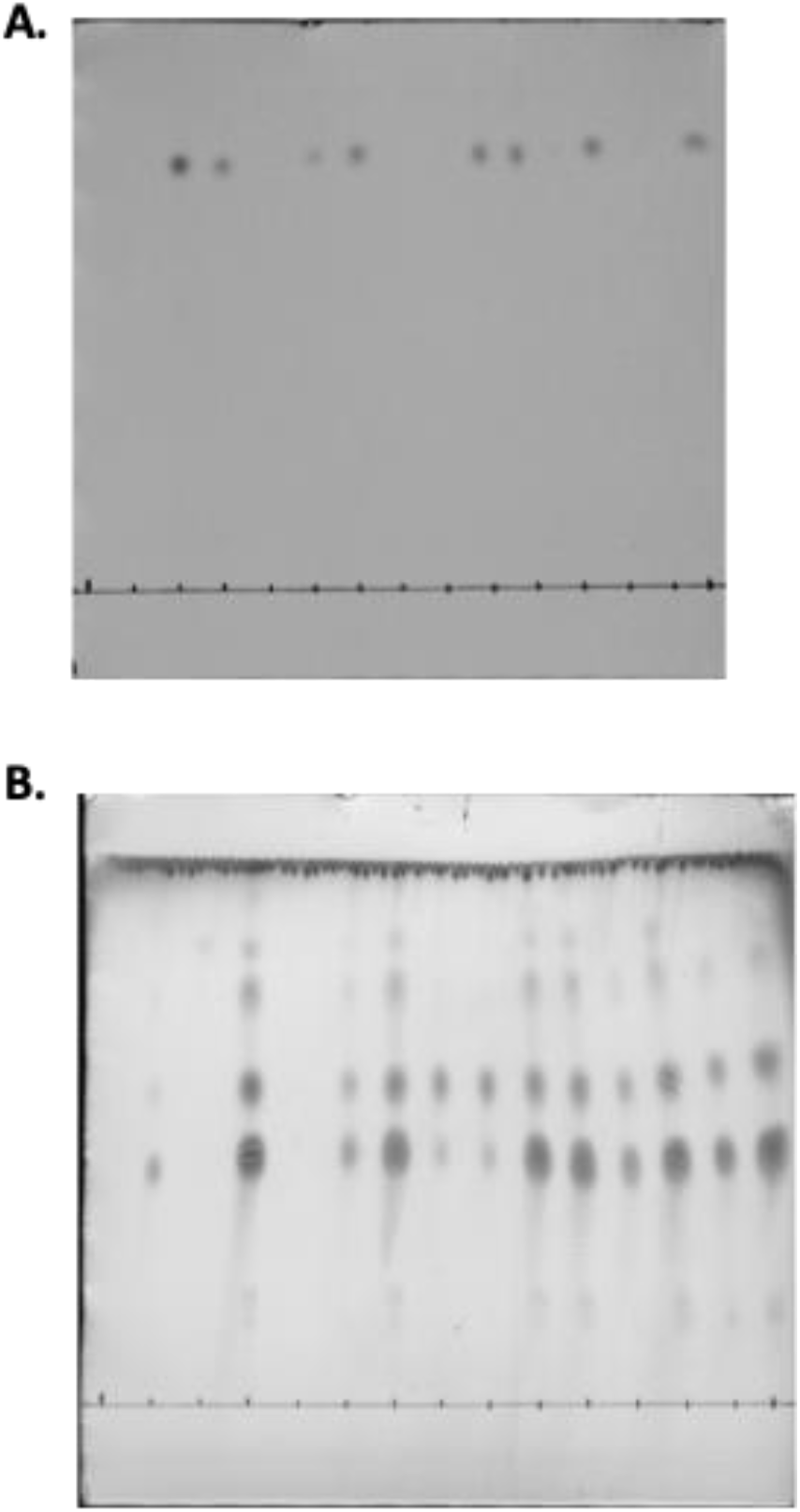
Raw images corresponding to figure 5. **A)** Complete image of TLC plate corresponding to figure 4B. **B)** Raw TLC image relating to figure 4C.

**Supplementary Figure 12.**
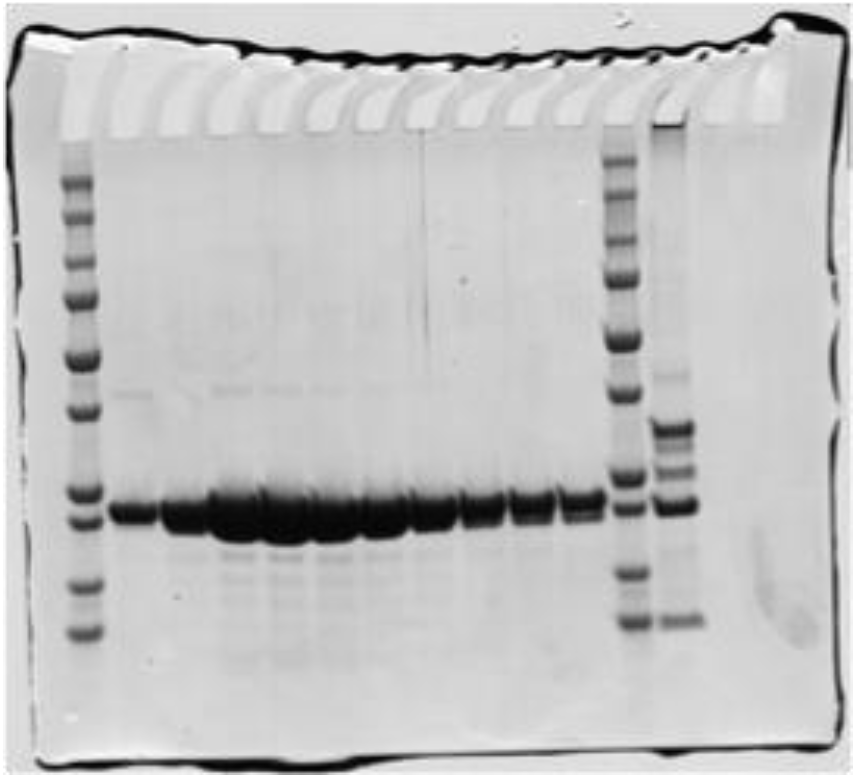
Raw image corresponding to supplementary figure 1A.

**Supplementary Figure 13.**
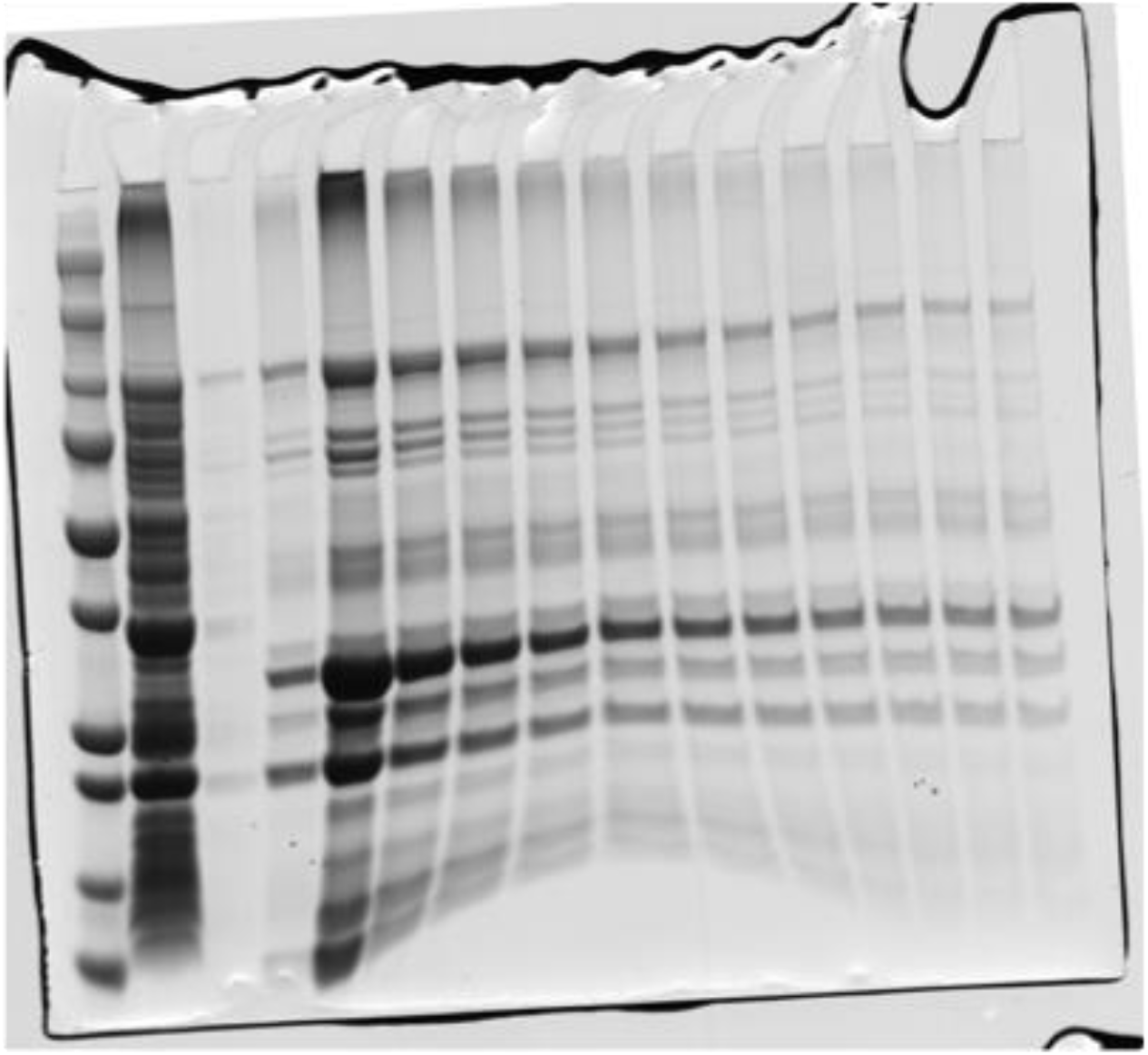
Raw image corresponding to supplementary figure 4A.

